# Expanding the Tubulin Code: TTLL11 Polyglutamylase Drives Elongation of Primary Tubulin Chains

**DOI:** 10.1101/2025.02.24.639257

**Authors:** Jana Campbell, Miroslava Vosahlikova, Samar Ismail, Margareta Volnikova, Lucia Motlova, Julia Kudlacova, Kseniya Ustinova, Ivan Snajdr, Zora Novakova, Miroslav Basta, Irina Gutsche, Marie-Jo Moutin, Ambroise Desfosses, Cyril Barinka

## Abstract

Microtubules (MTs) undergo diverse post-translational modifications that regulate their structural and functional properties. Among these, polyglutamylation – a dominant and conserved modification targeting the unstructured tubulin C-terminal tails – plays a pivotal role in defining the tubulin code. Here, we uncovered a novel mechanism by which tubulin tyrosine ligase-like 11 (TTLL11) expands and diversifies the code. Cryo-electron microscopy revealed a unique bipartite MT recognition strategy wherein TTLL11’s binding and catalytic domains engage adjacent MT protofilaments. Biochemical assays identified previously unknown polyglutamylation patterns, showing that TTLL11 directly extends the primary polypeptide chains of α- and β-tubulin, challenging the prevailing paradigms emphasizing lateral branching. Moreover, cell-based and *in vivo* data firmly established a crosstalk between TTLL11-mediated polyglutamylation and other tubulin-modifying processes, notably the detyrosination/tyrosination cycle. This discovery unveils an unrecognized layer of complexity within the tubulin code and offers new insights into the molecular basis of functional specialization of cytoskeleton across diverse cellular contexts.

## Introduction

The cytoskeleton serves as the structural scaffold and dynamic machinery orchestrating vital cellular processes, ranging from cell division to intracellular transport. Microtubules (MTs) represent fundamental constituents of this framework. MTs exhibit high genetic diversity arising from the existence of multiple isoforms of α- and β-tubulin that are encoded by distinct genes with tissue-specific expression patterns (*1*). Additionally, a plethora of post-translational modifications (PTMs) further diversifies the MT landscape, modulating their stability, interactions, and functional properties. Altogether, these variations comprise the so-called tubulin code (*2*).

Several PTMs, including acetylation, methylation, and phosphorylation, are confined to the tubulin structured core. However, in the context of the MT surface and interactions with molecular effectors, modifications to unstructured C-terminal tubulin tails might be far more impactful (Fig. 1A) (*3*). A genetically encoded C-terminal tyrosine of the α-tubulin primary polypeptide chain is removed by either the vasohibin/small vasohibin-binding peptide complex (VASH/SVBP) (*4, 5*) or tubulin metallocarboxypeptidase 1 (TMCP1) creating the αΔTyr variant (*6, 7*). The two remaining terminal glutamates can be further trimmed by the action of cytosolic carboxypeptidases (CCPs) (*8, 9*) or TMCPs (*7*) generating αΔ2 and αΔ3 tubulins. While the removal of the C-terminal tyrosine can be reversed by tubulin tyrosine ligase (TTL) (*10*), the deletion of the terminal glutamates is considered irreversible (*11*). Truncations of the primary polypeptide chain by TMCP2 have also been reported for several β-tubulin isoforms (*7, 12*). As in the case of αΔ2/αΔ3 tubulins, no rescue of the truncated β-chains has been reported in the literature (Fig. 1B).

**Fig. 1.**
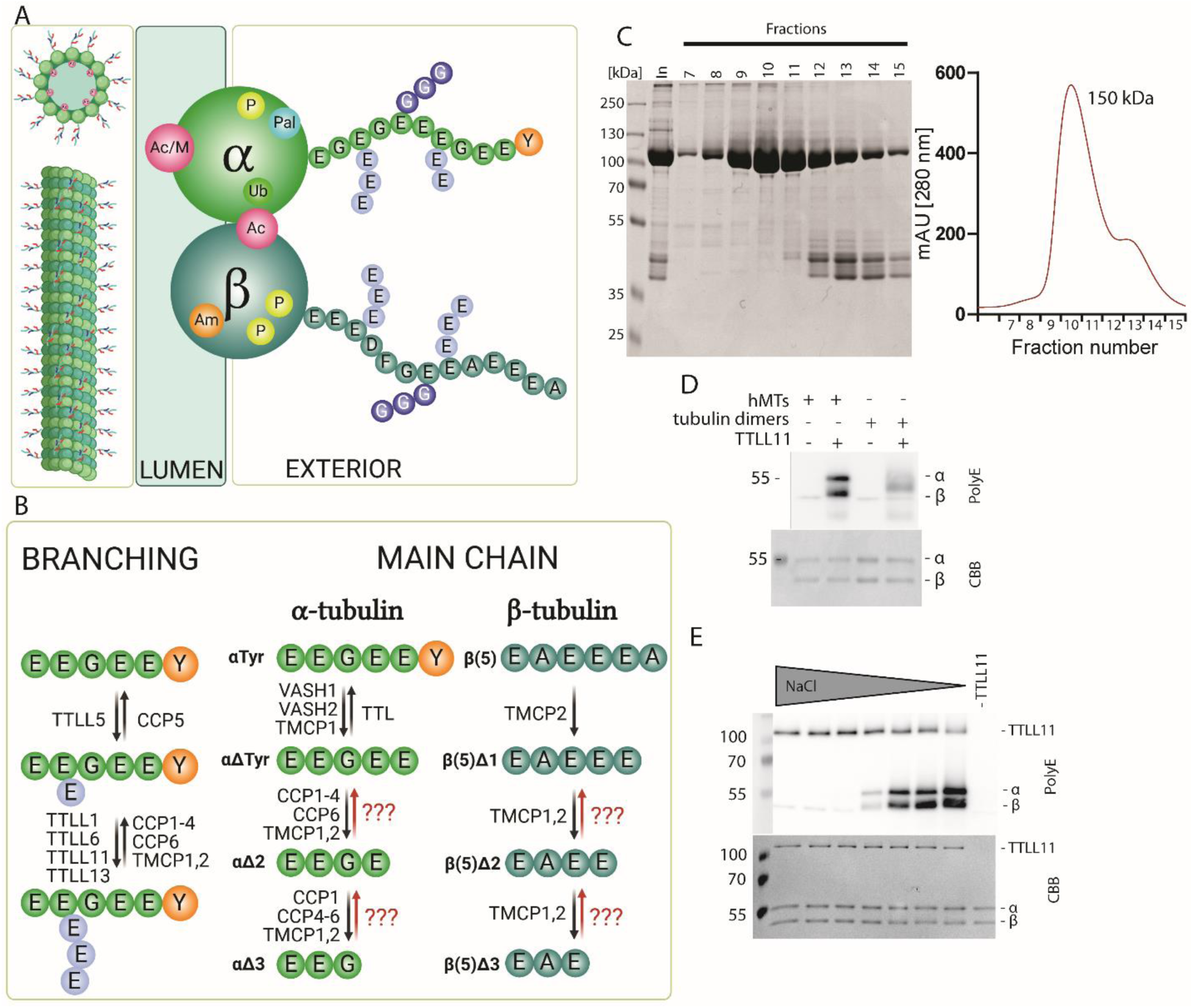
The tubulin code, TTLL11 purification and tubulin polyglutamylation. **A. Post-translational modifications defining the tubulin code** are found at structured tubulin cores (acetylation [Ac], methylation [M], phosphorylation [P], amination [Am], palmitoylation [Pal], ubiquitination [Ubl]) and at disordered C-terminal tails (glycylation [G], glutamylation [E]). **B. Glutamylases of the TTLL family catalyze branching and elongation** of polyE chains, which are removed by CCPs and TMCPs. The reversible detyrosination/tyrosination cycle is mediated by TTL, VASHs, and TMCP1. CCPs and/or TMCPs truncate the C-termini of tubulin primary chains, yielding irreversible “dead-end” tubulin variants. **C. TTLL11 purification.** Size-exclusion chromatogram and corresponding Coomassie Brilliant Blue (CBB) stained PAGE gel from purification of human TTLL11 expressed in HEK293T cells. The main peak corresponds to a monomeric TTLL11 species (∼150 kDa). **D. TTLL11 prefers microtubules (MTs) over tubulin dimers**, polyglutamylating both α- and β-tubulins. Western blot visualized with the polyE antibody (upper panel) together with the CBB-stained PAGE gel as protein loading control (lower panel). **E. TTLL11 glutamylation activity correlates inversely with ionic strength.** TTLL11 efficiently polyglutamylates MTs in low ionic strength buffers (BRB30 + 1 mM NaCl) with reduced activity observed in increasing salt concentrations. The WB upper ∼120 kDa band represents autoglutamylated TTLL11. The CCB-stained gel as a protein loading control.

In addition to primary chain truncations, α- and β-tubulin tails can be laterally branched by polyglutamate chains attached to the γ-carboxyl group of internal glutamate residues (Fig.S1). Glutamylation is evolutionarily conserved from ciliates to humans; it is a hallmark of differentiated cells, such as neurons, and is predominantly found in tubulin-rich substructures including axons, dendrites, cilia, flagella as well as the mitotic spindle. Expectedly, glutamylation homeostasis is tightly regulated as it is critical for normal cell physiology and aberrant polyglutamylation is linked to neurodegeneration (*8*), ciliopathies (*13, 14*), immune response defects (*15*), and cancers (*16–18*).

Two principal glutamylation steps, branching and elongation, are catalyzed by several members of the tubulin tyrosine ligase-like (TTLL) family, each with distinct substrate specificities (Fig. 1B) (*19*). While tubulin is the best-studied TTLL substrate, polyglutamylation has been reported to modulate the physiological functions of other protein targets such as Dishevelled, cyclic GMP-AMP synthase, and histone chaperones (*15, 20, 21*). Moreover, existing data suggest that glutamylation might be more widespread than currently acknowledged and the TTLL11 enzyme can play a prominent role in polyglutamylation of non-tubulin substrates (*22*). Depending on the organism and the cell type, TTLL11 is localized in the nucleus in human fibroblasts and HeLa cells (*23*), but it was observed in the cilium in *C. elegans* (*24*) and the basal body of MDCK cells (*19*). TTLL11 is critical for glutamylation of the mitotic spindle, and its silencing decreases the chromosome segregation fidelity associated with cancer (*18*). At the organism level, TTLL11 mutations affect skeletal development in humans and these observations were replicated in zebra fish, where TTLL11 mutations led to mostly fatal spine curvature defects (*25*).

By combining cryo-EM and functional assays, this study uncovers a novel type of tubulin glutamylation pattern, where TTLL11 extends the primary polypeptide chains of both α- and β- tubulin with specificity driven by the amino acid composition of the respective C-terminal residue. It further reveals a crosstalk between other tubulin modifying enzymes and TTLL11, and specifically between the vital detyrosination/tyrosination cycle of α-tubulin and TTLL11 polyglutamylase. This discovery challenges the existing understanding of tubulin modifications, which primarily emphasized lateral branching polyglutamylation. Our work thus significantly advances the field by expanding the known repertoire of modifications within the tubulin code.

## Results

### TTLL11 polyglutamylates microtubules at both α- and β-tubulin chains

To elucidate the substrate specificity of TTLL11, we expressed and purified the human enzyme (human TTLL11; Fig. 1C) and assayed its competency to modify tubulins isolated from HEK293T cells. Our data clearly show that TTLL11 preferentially polyglutamylates polymerized MTs over free tubulin (Fig. 1D) in an ionic strength-dependent manner (Fig. 1E; Fig. S2A-C). Interestingly, our findings also reveal that both α- and β-subunits are polyglutamylated by hTTLL11 *in vitro*, indicating that the enzyme exhibits a broad substrate specificity that was not previously recognized (*19, 22*). The lack of hTTLL11 selectivity for either α- or β-tubulin thus contrasts with the substrate specificity of TTLL6 and TTLL7 paralogs with ascribed preferences for α- and β-chains, respectively (*26, 27*).

### TTLL11 simultaneously engages adjacent microtubule protofilaments

To provide the structural basis for TTLL11 substrate recognition and interpret potential preferences for polymerized MTs as well as α- vs β-tubulin chains, we determined the cryo-EM structure of hTTLL11 (residues 128 – 657) in the complex with double-stabilized MTs (dsMTs) isolated from HEK293T cells. The final cryo-EM map has a nominal resolution of 3.28 Å with local resolution estimate up to 2.8 Å (Fig. 2A-C; Fig. S3A-D). The well resolved map facilitated the unambiguous assignment of individual tubulin protomers (Fig. S3E) as well as the construction of a high-confidence atomic model comprising the newly identified MT-binding helix bundle of hTTLL11 (MT-BHB; residues C531 – R659; Fig. 2D,E). The MT-BHB, which does not have any homology counterparts in MT-binding motifs of other members of the TTLL family, represents a critical component of the TTLL11/MT interaction interface (Fig. 2F, Fig S3F,G) (*28, 29*). Although the density representing the TTLL11 catalytic core (residues P128 – P486) is less resolved, it was still possible to reliably dock the AlphaFold TTLL11 model into the cryo-EM map. The following segment (K488 - V535) connecting the catalytic domain to the MT-BHB is not visible in the map and therefore not included in the model, nor are the intrinsically disordered N- and C-termini, which are also missing from the cryo-EM map. Taken together, the cryo-EM data indicate that while a newly identified MT-BHB is rigidly docked onto the MT surface, the TTLL11 catalytic core (together with intrinsically disordered regions) and tubulin C-terminal tails are inherently flexible and can adopt a wide range of conformations.

**Fig. 2.**
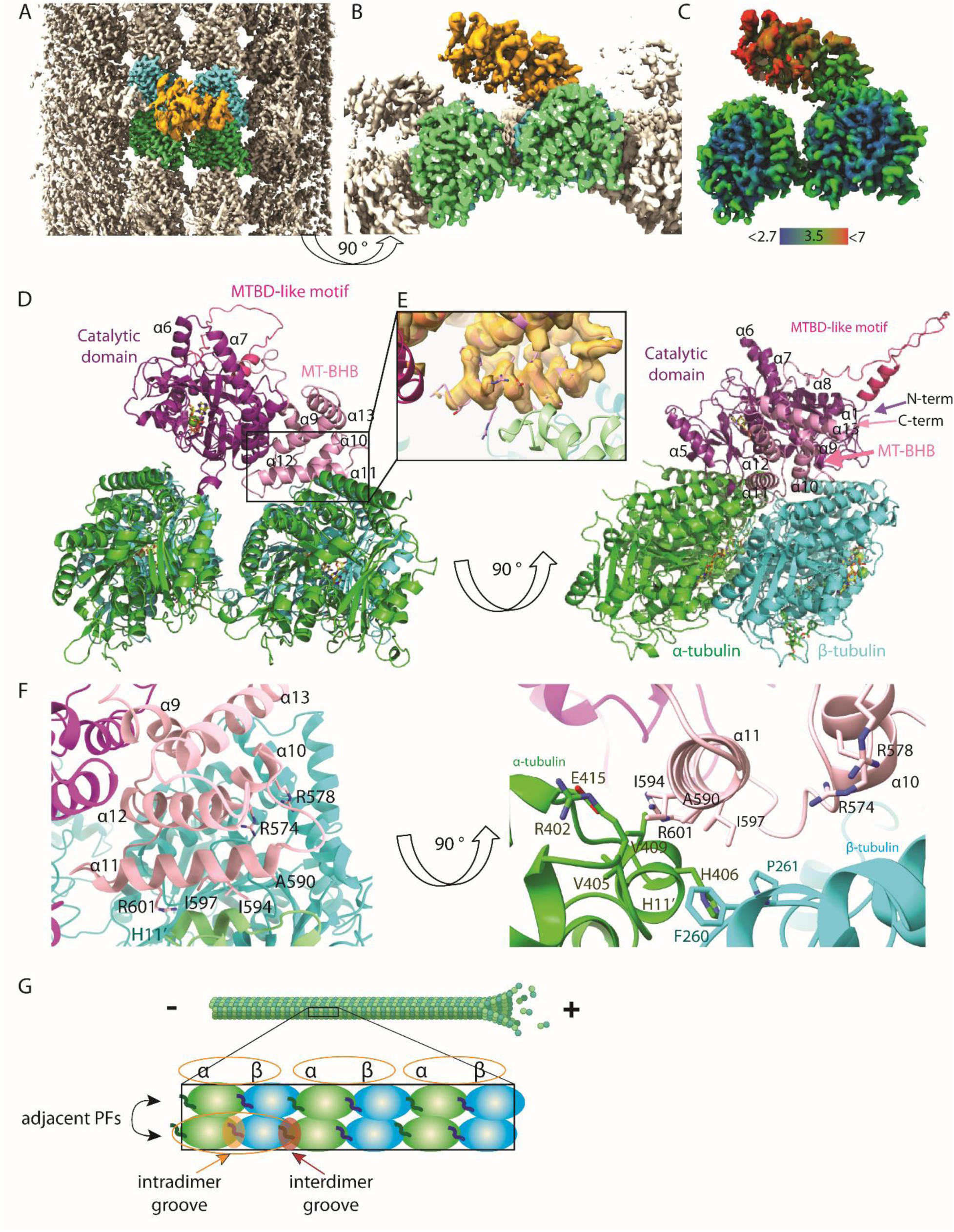
The cryo-EM reconstruction of the TTLL11/MT complex. **A, B. The cryo-EM map shows TTLL11 binding to the microtubule (MT) lattice**. TTLL11 in gold, MT in grey, and the central α- and β-tubulin dimers in green and cyan, respectively (**A**: top view; **B**: side view). **C. Cryo-EM map colored by local resolution**, ranging from 2.7 Å (blue, tubulin core) to over 7 Å (red, distal TTLL11 catalytic domain). **D. A cartoon model of the TTLL11/MT complex**. TTLL11 binds the α/β intradimer interface via its microtubule-binding helix bundle (MT-BHB, pink, 9HQ4), positioning the catalytic domain (magenta) over the β-tubulin C-tail of an adjacent protofilament. The helix-loop-helix motif (hot pink, MTDB-like) is structurally analogous to TTLL6/7 binding domains but is not implicated in MT interactions. **E, F. Details of the TTLL11/MT interface.** The detailed view of interactions between α11 helix of the MT-BHB modeled into the cryo-EM map (gold) and the MT surface **(E)**. Zoomed view of interface between MT-BHB and tubulin dimer (helix H11’ and loop between S8 and H10’ of α- and β-tubulin, respectively (*30*) (**F**). Residues A590, I594, I597, and R601 (helix α11), and R574/R578 (helix α10) of TTLL11 contribute prominently to the mixed ionic/hydrophobic interface with MTs. **G. Schematic representation of MT protofilaments**. MTs are polar tubular assemblies of α/β-tubulin dimers, with β-tubulin on the (plus)-end. Dimers form protofilaments (PFs), stacked side-by-side in a circular arrangement. Intradimer grooves (yellow ovals) are located between α- and β-tubulin in dimers, while interdimer grooves (red ovals) separate adjacent dimers.

Two unexpected findings emerged from our cryo-EM reconstruction. First, TTLL11 recognizes MTs in a unique bipartite pattern, where the MT-BHB and the catalytic domain engage adjacent protofilaments (PF). This stands in stark contrast with the structures of TTLL6 and TTLL7 complexes, where both MT binding and polyglutamylation are confined to the same PF (Fig. S4) (*26, 29*). Secondly, the newly identified MT-BHB interacts with residues of the longitudinal intradimer α/β groove, positioning thus the catalytic TTLL11 domain atop the C-terminal tail of the β-subunit of the adjacent PF (Fig. 2F,G; Fig S3H-J). These findings are consistent with the biochemical data showing that the tubulin tetramer within the MT lattice, but not the tubulin dimer, represents the minimum TTLL11 recognition motif (Fig. 1D). At the same time, positioning of the catalytic domain above the β-subunit C-terminal tail would point towards a preference for polyglutamylation of β-tubulin. However, both α- and β-tubulin chains of the tubulin isolated from HEK293T cells are polyglutamylated to a similar extent *in vitro* (Fig. 1D,E).

### The MT-BHB does not discriminate between intra- and inter-dimer tubulin interfaces

Sequence alignment and structural models clearly indicate that the MT-BHB of TTLL11 is distinct from known or presumed MT-binding segments of other TTLL family members (Fig. S5; S6). The structured part of the MT-BHB forms a five-helix bundle (comprising helices α9 through α13) that is in apposition to the catalytic domain with a shared interface of approximately 1500 Å^2^. The cryo-EM structure revealed that the primary MT interaction motif of MT-BHB involves two amphipathic helices encompassing amino acids T571 – C580 (helix α10) and M589 – R601 (helix α11; Fig. 2E,F, Fig. S5D). The positively charged R574 and R578 (α10) and R601 (α11) are oriented towards the negatively charged tubulin surface to mediate putative ionic interactions. However, the key interaction motif involves the hydrophobic face of the α11 helix comprising A590, I594, and I597 that is engaged by residues of the H11’ helix of α-tubulin and the side chains of F260/P261 of the β-subunit (Fig. 2F) (*30*). The pivotal contribution of the side chain of I594, which is inserted into the complementary tubulin hydrophobic pocket delineated by the side chains of R402, V405, H406, V409, and E415 (Fig. 2F; Fig. S3F,G; Fig. S7A,B), was corroborated by side-directed mutagenesis and *in vitro* experiments (see below).

Given the promiscuity of TTLL11 towards α- and β-tubulin tails, we further analyzed the sequence and structural conservation of tubulin residues implicated in interactions with MT-BHB. Overall, the high sequence/structural conservation between individual tubulin isoforms as well as between α- and β-tubulins (Fig. S7A,B) indicates that while MT-BHB is conceivably the primary MT recognition motif anchoring TTLL11 to the MT surface, it alone is unable to discriminate between intra- and interdimer groove interfaces of tubulin within the MT lattice. Consequently, isolated MT-BHB cannot dictate TTLL11 preferences for either α- or β-protomer and TTLL11 selectivity (or lack thereof) for either tubulin protomer requires additional inputs from the catalytic domain.

### Bipartite engagement is essential for efficient MT binding and polyglutamylation by TTLL11

To evaluate if and how the interplay between the catalytic domain and MT-BHB influences TTLL11 interactions with MTs, we generated a series of TTLL11 variants. These variants were assayed for their ability to bind MTs using total internal reflection fluorescence (TIRF) microscopy and modify MTs using the polyglutamylation assay. For the TIRF binding assay, porcine dsMTs were immobilized onto coverslips and the binding of fluorescently labeled TTLL11 variants quantified. While the strong fluorescence signal was observed for the full length TTLL11 as well as for variants lacking the N-terminal region (M1 - G121), neither MT-BHB nor the catalytic domain was able to bind MTs in isolation (Fig. 3A,B). Furthermore, the importance of MT-BHB (and its crosstalk with the catalytic domain) was underscored by the analysis of the I594W and R601E mutants, which were designed to impair interactions with MTs based on our cryo-EM structure. Here, the fluorescence signal intensity was reduced by 60 % and 80 % for the R601E and I594W variant, respectively. Similarly to the glutamylase activity, the TTLL11 binding to MT surface was also dependent on the ionic strength of the assay buffer (Fig. S2).

**Fig. 3.**
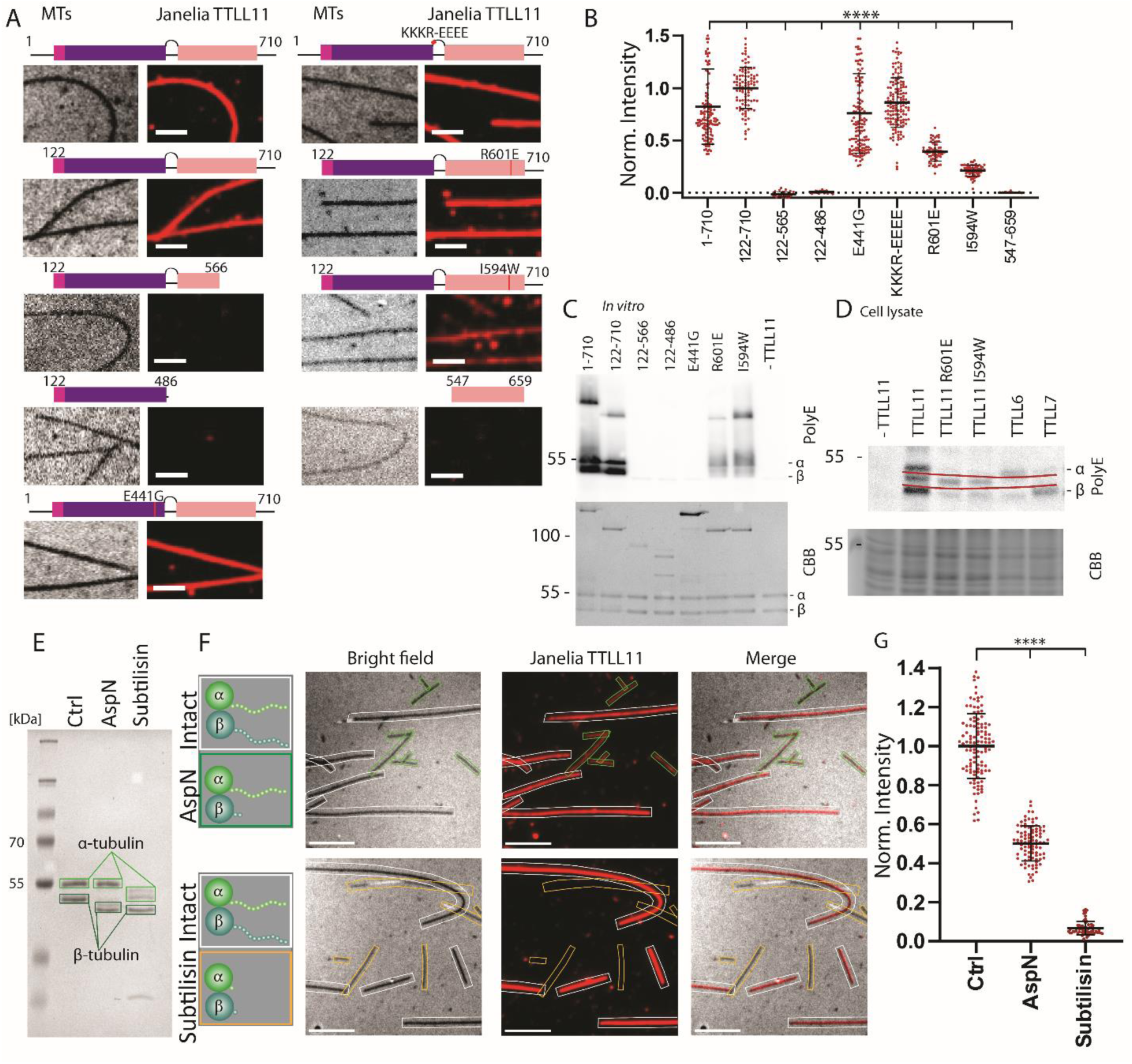
Crosstalk between the catalytic and MT-BHB domains is critical for MT-binding and polyglutamylation by TTLL11. **A. TTLL11 binding to unlabeled porcine MTs analyzed using TIRF microscopy**. MTs were attached to coverslips and incubated with 100 nM Janelia549-labeled TTLL11 variants (red). Scale bar: 2 µm. **B. Quantification of TIRF images** for full-length TTLL11, the N-terminally truncated variant (122–710), the catalytically inactive E441G mutant, and the KKKR-EEEE mutant show binding with similar intensity. Deleting the MT-BHB or catalytic domains abolishes TTLL11 binding. Mutations in TTLL11 helix α11 residues (R601E, I594W) reduced binding by 60% and 95%, respectively. C. *In vitro* polyglutamylation of purified human MTs by TTLL11 variants, analyzed by Western blotting, confirms robust activity for full-length and 122-710 variants. Variants lacking the MT-BHB domain show no activity. I594W and R601E mutants show reduced activity due to weakened MT interactions. D. *In-cellulo* assays in HEK293T cells transfected with TTLL11 variants, TTLL6, and TTLL7 have polyglutamylation patterns consistent with *in vitro* findings. TTLL6 and TTLL7 show specificity for α- and β-tubulin, respectively. An unidentified TTLL11 substrate, represented by a polyE-stained band in between α- and β-tubulin (marked by red lines), is glutamylated regardless of the MT-BHB mutations. **E - G. Tubulin C-tails are critical for TTLL11 interactions. E.** CBB-stained gel of AspN and subtilisin-treated MTs. AspN truncates the β-tubulin C-tail, while subtilisin truncates both α- and β-tubulin C-tails. TIRF microscopy images (**F**) and quantification (**G**) show AspN-treated MTs exhibit ∼50% reduction in TTLL11 binding, while the complete loss of binding is observed for subtilisin-treated MTs. Data is shown as mean fluorescent intensity from n = 2 replicates with 114, 93, 58 MTs quantified in each sample. Statistical significance was determined using the unpaired t-test with Welsch correction, ****p<0.0001, the black bar represents median value with 95 % c.i. Scale bar = 5 µm.

In a complementary set of experiments, we generated modified MTs in which the C-tails of dsMTs were removed by treatment with AspN (β-tail only) or subtilisin (both α- and β-tails; Fig. 3E). At 100 nM concentration, the fluorescence TTLL11 signal for AspN-treated MTs was approximately 50 % lower compared to intact MTs, and no binding was observed for subtilisin-treated MTs. Clearly, the C-terminal tail of both tubulin protomers is essential for anchoring TTLL11 to MTs and the monodentate binding mediated by MT-BHB is not sufficient. The combined contribution of both TTLL11 domains is thus essential for effective MT binding (Fig. 3F,G).

The TIRF experiments were further corroborated *in vitro* and cell-based assays where we evaluated MT polyglutamylation by TTLL11 variants (Fig. 3C,D). While wild type TTLL11 showed pronounced tubulin polyglutamylation, the presence of the inactive E441G mutant or the isolated catalytic domain did not change polyglutamylation beyond background levels. Similarly, tubulin polyglutamylation was markedly impaired in the case of the I594W and R601E mutants. Combined, these results show that the lower binding affinity of TTLL11 variants is directly translated into less efficient polyglutamylation of the target tubulin substrate.

### TTLL11 expands the tubulin code by extending the primary tubulin chains

While Western blotting is a valuable tool for qualitative analysis of polyglutamylation in cells and *in vitro*, it does not allow for the identification of polyglutamylation attachment sites and polyglutamate chain length and connectivity. To overcome these limitations, we implemented an LC-MS/MS pipeline that enabled us to gain detailed qualitative and quantitative insights into tubulin polyglutamylation by TTLL11. For MS/MS experiments, samples were digested with the trypsin/LysC mix or AspN for the analysis of the C-terminus of α- and β-tubulin, respectively. MS data were used for quantification, while MS/MS spectra provided information on the positions of lateral branching points and linkage chemistry. Accurate identification of fragment peaks was facilitated by exploiting isotopically labeled perdeuterated-D_5_ and ^18^O glutamates (Fig. 4A).

**Fig. 4.**
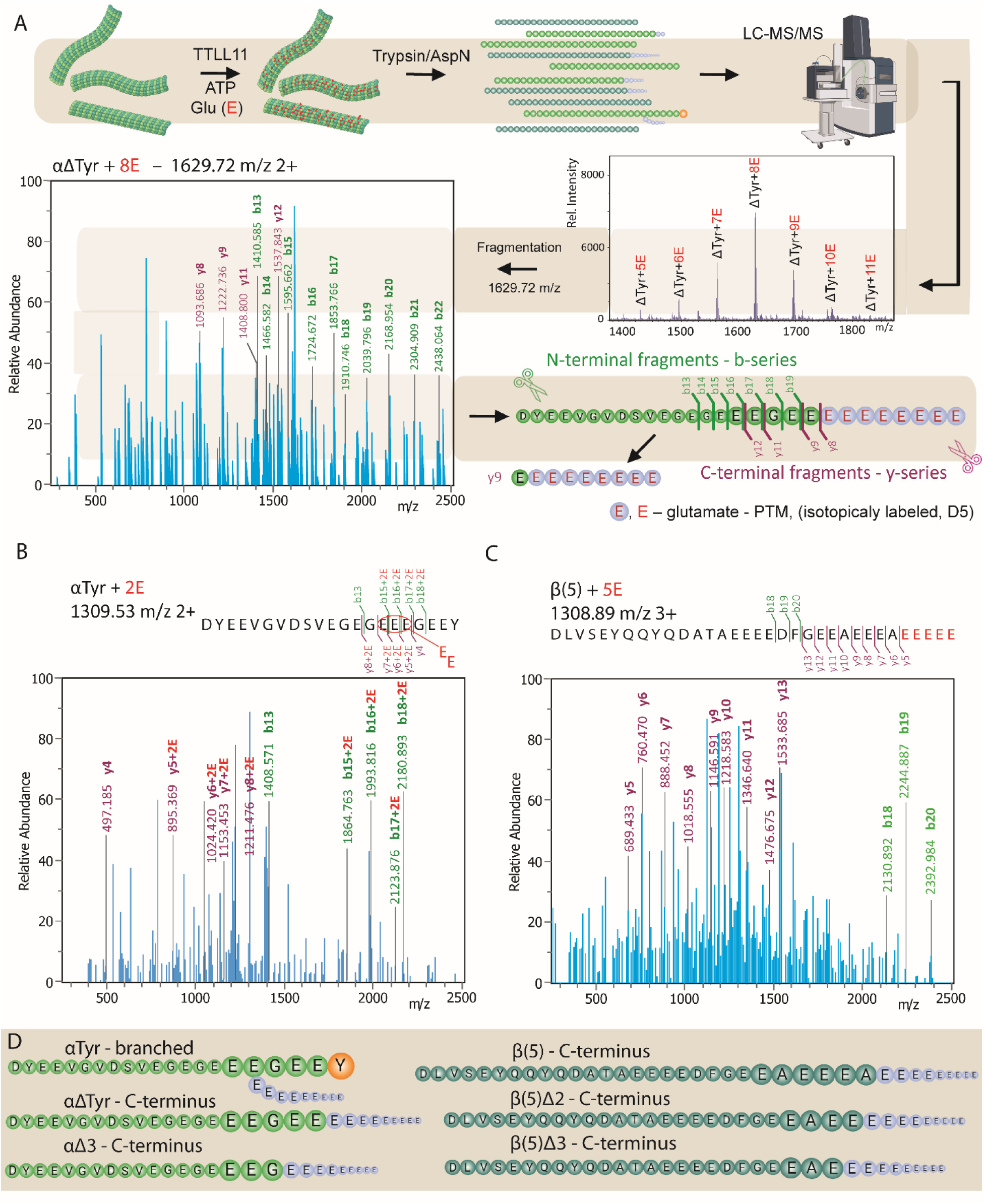
Identification of polyglutamylation sites within tubulin sequences using the LC-MS/MS pipeline. **A. Schematic representation of the LC-MS/MS pipeline**. MTs are incubated with TTLL11 *in vitro* in a reaction mixture supplemented with perdeuterated D5-glutamate. Following digestion with either trypsin (α-tubulin C-tails) or AspN (β-tubulin C-tails), resulting peptides are analyzed by LC-MS/MS for the presence of (poly)glutamylated species. For each of the (poly)glutamylated peptides, MS/MS fragmentation is performed to identify the site(s) of attachment of the polyE chain(s), based on the presence of the b- and y-series of peptides, which correspond to N- and C-terminal fragments, respectively. **B, C. Illustrative examples of MS/MS spectra of polyglutamylated tubulin species.** The LC-MS/MS pipeline was used to analyze tubulin variants that were polyglutamylated by TTLL11 *in vitro*. Peaks of interest observed in b- and y-series are highlighted in green and magenta, respectively. This approach enabled the unequivocal assignment of attachment sites polyglutamate chains within the tubulin sequence. **D. Summary of tubulin sites preferentially polyglutamylated by TTLL11.** The combined LC-MS/MS data reveals that the αTyr variant is branched (and extended) at positions E445, E446, and E447, albeit at marginal levels. In contrast, the remaining tubulin variants are preferentially polyglutamylated at their C-termini by the direct extension of the primary polypeptide chain. The corresponding MS and MS/MS spectra and quantifications are shown as Supplementary Figures S9 through S12.

In our original set of experiments, we used mostly unmodified native tubulin isolated from HEK293T cells, where TUBA1A and TUBA1B isoforms, sharing identical C-terminal sequences, account for over 80 % of all α-tubulins (Fig. S7C,D; S8) (*31*). Furthermore, tyrosinated α-tubulin (αTyr) represents approximately 80%, while αΔTyr and αΔ2 variants are much less populated (approximately 15%). The remaining 5% minority species included C-tails with one or two glutamate residues added.

Upon incubation with purified TTLL11, we observed massive polyglutamylation of predominantly αΔTyr and αΔ2 variants with up to 11 glutamate residues added. Strikingly, the MS/MS fragmentation spectra revealed that the glutamate residues are primarily attached to the very C-terminus of the main polypeptide, comprising thus a new type of tubulin modification (Fig. 4A, Fig, S9A). The αTyr variant was not C-terminally extended but rather laterally branched at Glu445, Glu 446 or Glu447, albeit at marginal levels (Fig. 4B).

For β-tubulins, TUBB5 and TUBB4B were the most populated isoforms (approximately 80 %), followed by lower amounts of TUBB2A/B (Fig. S7D) (*31*). Less than 5 % of native β-tubulins were modified by an extra glutamate residue. When incubated with TTLL11 *in vitro*, up to 27 glutamate residues were attached to β-tubulin tails and the MS/MS spectra again revealed the direct extension of the native C-termini, where polyglutamate chains were attached to the terminal alanine residue shared by the TUBB5/TUBB4B isoforms (Fig. 4C,D; Fig. S9B,C).

### Main chain tubulin polyglutamylation is preferred over elongation of laterally branched chains

To analyze TTLL11 preferences for elongating of existing laterally branched chains versus extending the tubulin main chain, we used tubulins isolated from porcine brain as substrates for *in vitro* reconstitution assays. In contrast to unmodified HEK293T tubulin, porcine tubulin contains a diverse spectrum of modified tubulins with mono-and polyglutamylated species representing approximately 60% of the total α-tubulins. In addition, the αΔTyr and αΔ2 variants are also more abundant, accounting for approximately 60 % and 20 % of α-chains, respectively (Fig. S8D). After incubation with purified TTLL11, MS/MS analysis revealed the presence of tubulin species directly elongated at the C-terminus of αΔTyr as well as elongation at preexisting branching points. While the complexity of the substrate and product species in the reaction mixture did not allow for their precise quantification, MS/MS fragment intensities corresponding to the C-terminally extended variants were generally stronger compared to the branching point elongation (Fig. S9D-F), pointing towards a preference of TTLL11 for main chain elongation.

The above findings raised an interesting idea that other elongases may also be capable of extending the primary tubulin chain, either independently or in conjunction with the elongation of the preexisting glutamate lateral branches (*27*). To test this hypothesis, we employed MS/MS to analyze tubulin polyglutamylation by TTLL6 *in vitro* which preference for αΔTyr is known (*27*). Our data demonstrate that TTLL6 is capable of elongating both the branching points as well as the main chain of the αΔTyr variant (Fig. S10A,C,D). In accordance with the known TTLL6 preferences, marginal elongation of the C-termini of β-tubulins was observed (Fig. S10B). Surprisingly, direct extension of the tubulin main chain may thus represent a common enzymatic activity of TTLL elongases, a possibility raised by assessing TTLL6 substrate preferences at the peptide level (*27*) but never reported and studied in the context of MTs, as the major physiological target of TTLLs.

Taken together, these data conclusively demonstrate that TTLL11 elongates both α- and β-tubulin main chains, thereby expanding a portfolio of modifications that comprise the tubulin code. While TTLL11 can in principle also laterally branch the internal C-tail glutamates of α-tubulins, this reaction is extremely inefficient, and we believe it might simply be an artifact of *in vitro* reaction conditions (Fig. 5A,B; Fig S11A,B). Indeed, when analyzing the tubulin polyglutamylation patterns in reactions utilizing ^18^O-labeled glutamate, the MS spectra were dominated by peptide envelopes retaining the ^18^O isotope, thus providing further experimental evidence that the new peptide bonds are formed via the α-carboxyl group linkage rather than γ-carboxylate lateral branching (Fig. S10E,F). Therefore, we propose that the enzyme acts exclusively as an elongase under physiological conditions (*27*) corroborating its proposed functions in an earlier study done in HeLa cells (*19*).

**Fig. 5.**
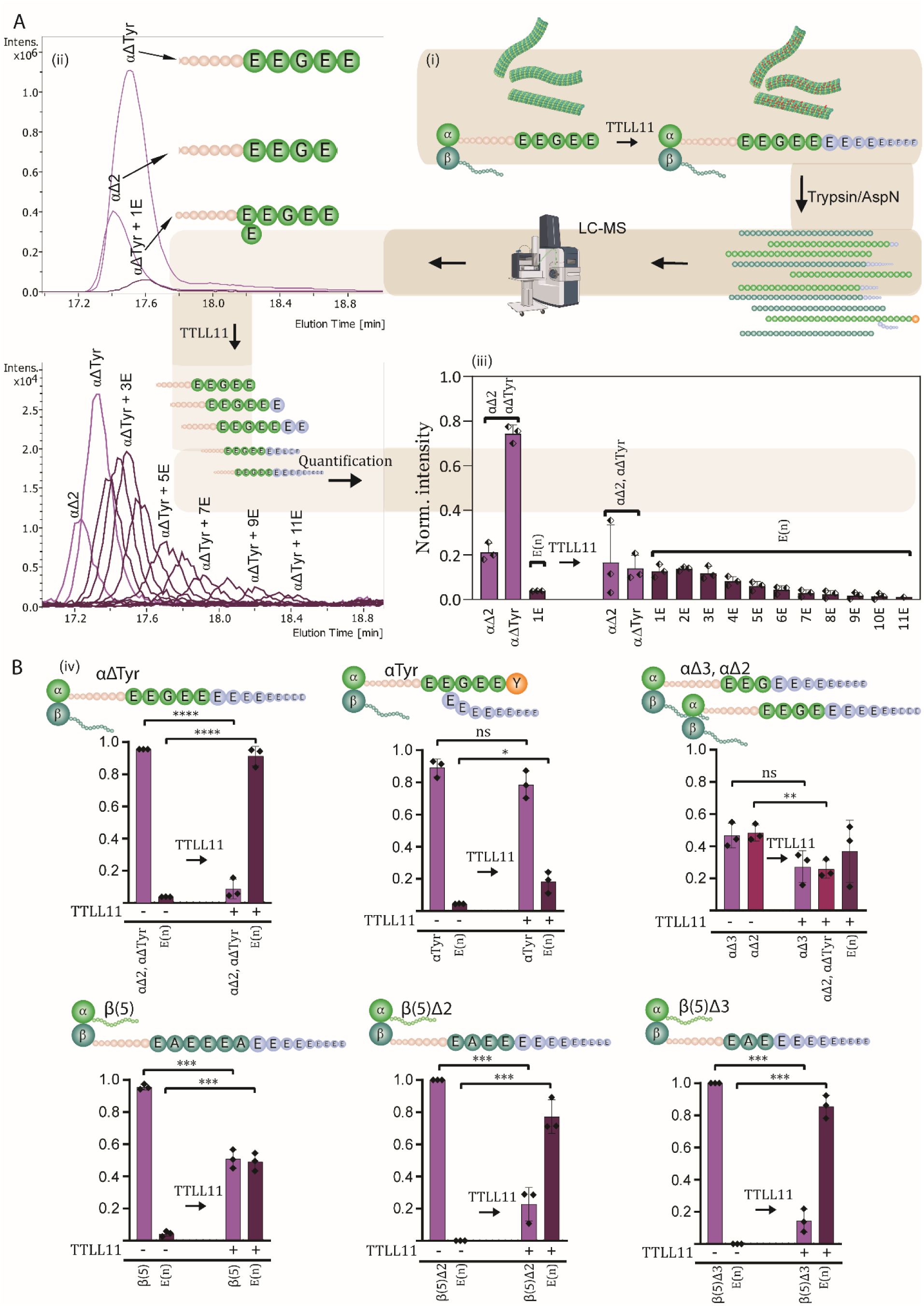
Quantitative analysis of tubulin polyglutamylation (polyE) by TTLL11. **A. An illustrative example of the quantification of tubulin C-tail polyglutamylation by LC-MS.** (i) Sequences of the C-terminal tryptic peptide of detyrosinated α-tubulin (αΔTyr) before and after TTLL11-mediated polyglutamylation. (ii) The EIC chromatograms show peaks corresponding to the peptide without glutamylation and with 0 – 11 glutamate residues attached. (iii) Quantification of the individual C-tail peptides with different polyE chains. (iv) A summary graph showing the normalized MS intensities before and after TTLL11 treatment derived from (ii) and (iii). **B. Summary quantification of tubulin polyglutamylation by TTLL11**. Tubulin variants (in the form of MTs) were incubated with TTLL11 and intensities of peptides before and after reaction were summed. The data are shown as the normalized MS intensities, where the sum of intensities of the peptide pool before and after TTLL11 treatment equals 1 (n = 3, statistical significance was determined using unpaired t-test, *p<0.05, **p<0.01, ***p<0.001, ****p<0.0001 ). While only marginal branched polyglutamylation is observed for the αTyr variant, remaining variants are efficiently polyglutamylated with conversions over 50% of the original tubulin (>80 % and >90 % for β(5)Δ2/β(5)Δ3 and αΔTyr, respectively). Detailed quantification of individual glutamylated peptides is shown in Fig. S11.

### Sequences of C-terminal tails dictate TTLL11 preferences for α- vs β-tubulins

Our data reveal that TTLL11 can extend the C-terminus of both α- and β-tubulins and that the C-terminal glutamate is not strictly required for the substrate to be modified. At the same time, when comparing the efficacy of polyglutamylation of αTyr vs αΔTyr (Fig. 5A,B; Fig. S11A,B), it is evident that TTLL11 discriminates between amino acid sequences of target substrates. To quantitatively analyze TTLL11 preferences for naturally occurring tubulin variants with different C-termini, we treated HEK293T tubulin with recombinant tubulin-modifying enzymes, yielding αTyr, αΔTyr, αΔ2, αΔ3, βΔ2, and βΔ3 enriched tubulin fractions for *in vitro* reconstitution experiments (Fig. S7C; S8A-C).

For α-tubulin variants, the main chain of either αΔTyr, αΔ2, or αΔ3 variants was directly extended with the reaction efficiency decreasing in the order of αΔTyr, αΔ2 > αΔ3, with 90% and 20% of the substrate modified, respectively (Fig. 5B, Fig. S11A-C). These results are particularly exciting with respect to the αΔ2 and αΔ3 variants, which were believed to be the dead-end products of α- tubulin C-tail modifications that could not be reverted to the native genetically encoded sequences (*11, 12*). Clearly, TTLL11 can rescue αΔ2 and αΔ3 variants forming αΔTyr that could either be tyrosinated by TTL or generate a novel type of C-terminal polyglutamate chain (Fig. 4D, Fig. S9A).

Recently identified TMCP enzymes were reported to remove residues from the C-terminus of β- tubulin, generating thus βΔ2 and βΔ3 variants (*7*). While native β-tubulins harboring the C-terminal alanine were glutamylated to approximately 50 %, the polyglutamylation efficacy of βΔ2 and βΔ3 variants was almost 80 % and thus comparable to results observed for αΔTyr (Fig. 5B, Figs. S8E, S11D-G). The C-terminal amino acid thus serves as the key determinant of TTLL11’s glutamylation activity and can determine TTLL11 preference for either α- or β-protomers (Fig. 6).

**Fig. 6.**
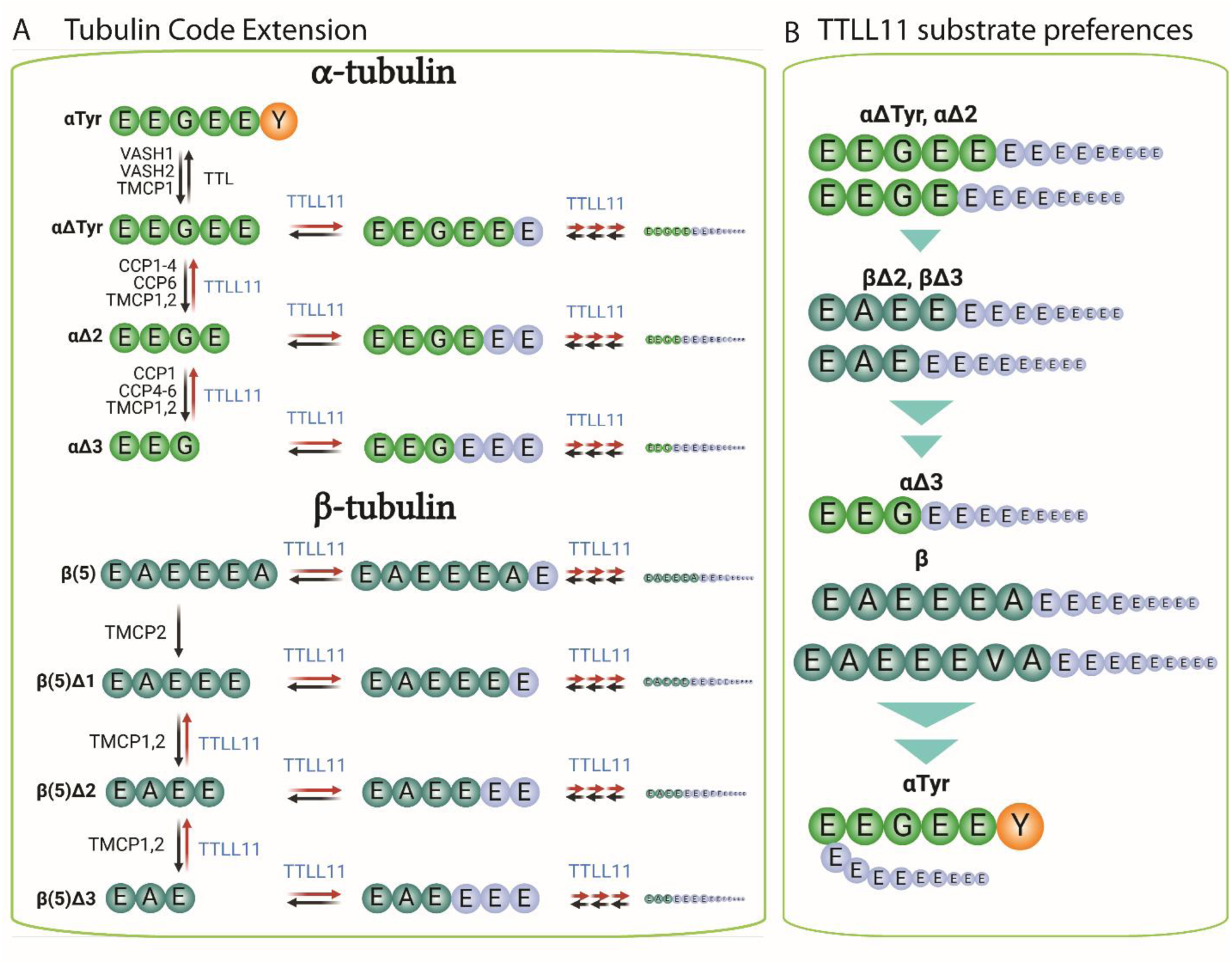
Expansion of the tubulin code by TTLL11-mediated polyglutamylation. **A. TTLL11 activity expands the tubulin code.** Polyglutamylation at the C-termini of the tubulin primary chains can rescue and recycle truncated tubulin variants that were previously considered irreversible end-products. Additionally, the main chain extension creates the new tubulin variants that could potentially be associated with hitherto unidentified physiological functions. **B. An overview of TTLL11 substrate specificity.** TTLL11 preferentially elongates the main chain of α- and β-tubulins with selectivity defined by the characteristics of the terminal amino acid. The ultimate glutamate residues are favored indicating that αΔTyr, αΔ2, βΔ2, and βΔ3 are optimal physiological substrates. As small aliphatic and hydrophobic residues are also permitted, αΔ3 and intact β-tubulins are also polyglutamylated with acceptable efficacy. The presence of the C-terminal Tyr/Phe prevents direct extension of the α-tubulin primary chain.

### Crosstalk between tubulin modifying enzymes and TTLL11 in cells and *in vivo*

We first replicated the biochemical *in vitro* experiments at the cellular level. Upon transfection of HEK293T and A549 cell lines with wild-type TTLL11, we observed the increase in the polyE signal intensities (Fig. 7A,B) for both α- and β-tubulins. In line with the low abundance of the αΔTyr variant (HEK293T), the most preferred TTLL11 substrate, the signal for polyglutamylated α-tubulin was less prominent than β-tubulin. Importantly though, cell transfections with vasohibin- 2/small vasohibin binding protein (VASH2/SVBP) led to the enrichment of the αΔTyr variant and a subsequent marked increase of the polyE signal of α-tubulin was observed in cells cotransfected with the combination of TTLL11/VASH2/SVBP (Fig. 7A,C). Interestingly, we could not observe any enrichment of β-tubulin polyglutamylation upon cell cotransfection with the combination of TTLL11 and *Tetrahymena* TMCP (ttTMCP; Fig. 7C). It is plausible that while the emergence of the Δβ2 variants would increase TTLL11 polyglutamylation rate towards this substrate as observed *in vitro*, ttTMCP, which has much higher *in vitro* hydrolytic activity compared to human enzymes, would simultaneously catalyze the removal of newly added polyglutamate chain from β-tubulin chains, thus counterbalancing TTLL11 activity. Clearly, and not surprisingly, the rate of α/β- tubulin polyglutamylation in a given cell would depend on the spatiotemporal distribution and activity of each of the tubulin modifying enzymes and their mutual interplay.

**Fig. 7.**
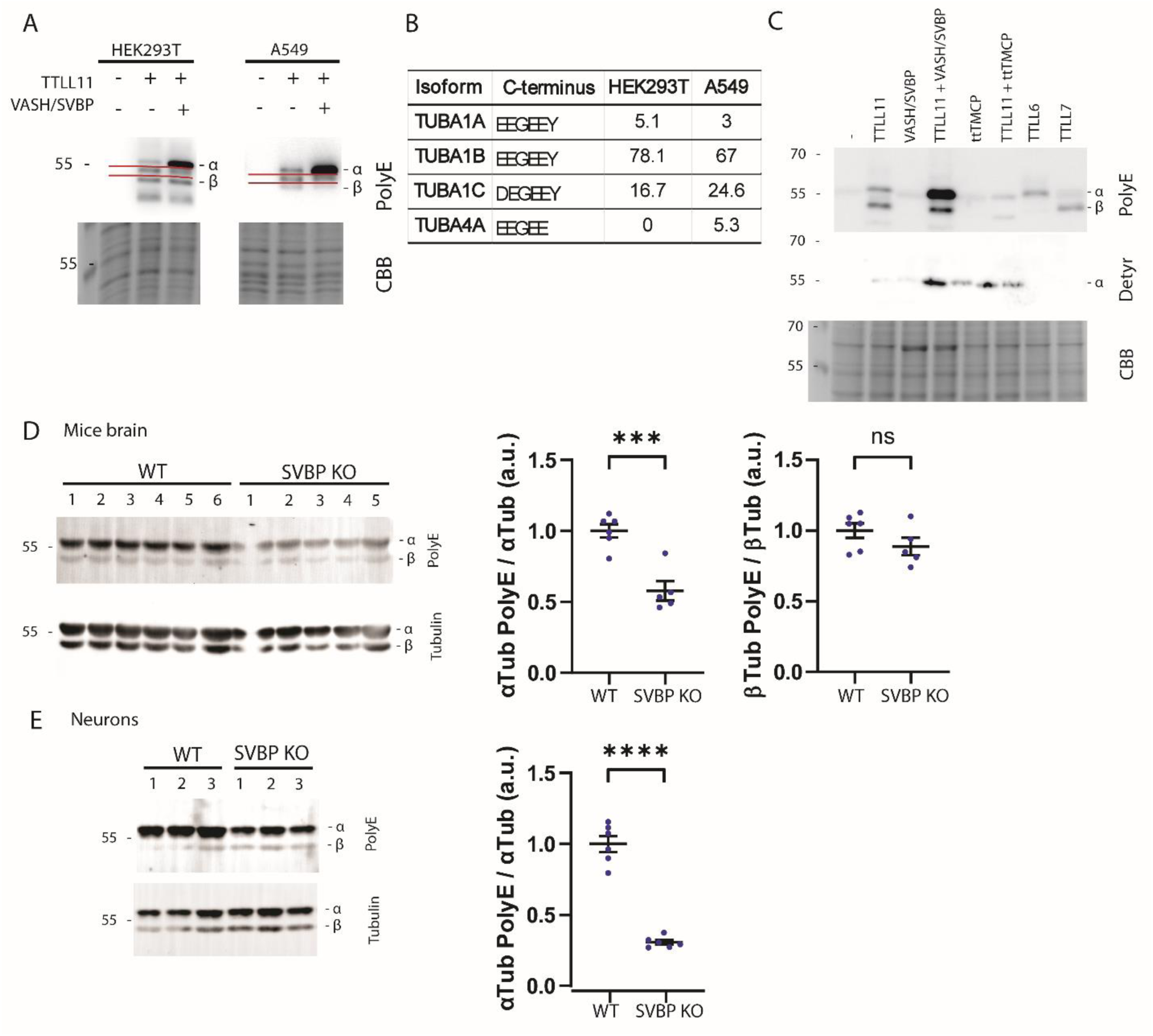
Crosstalk between TTLL11 polyglutamylase and tubulin modifying enzymes. **A-C. Protomer selectivity of TTLLs in cells.** WB (PolyE and deTyr antibodies - upper panel) of HEK293T/A549 cell lysates (co)-transfected with TTLL11, VASH2/SVBP, ttTMCP, TTLL6, and TTLL7. CBB-stained gel (lower panel) represents loading controls. **A. The ratio of α- vs β-tubulin polyglutamylation** depends in part on expression of the TUBA4A isotype. A549 cells have higher expression levels of the TUBA4A isoform, which lacks the C-terminal tyrosine and is a preferred TTLL11 substrate, leading to higher proportion of glutamylated α-tubulins. Red lines border a band of an unknown TTLL11 substrate. **B**. **Normalized mRNA transcript abundance** of α-tubulin isoforms in HEK293T and A549 cells was taken from (*31*). The TUBA4A isoform is present in A549 but absent from HEK293 cells. **C. VASH/SVBP transfection increases polyglutamylation of α-tubulin**. α-tubulin polyglutamylation of cells co-transfected with TTLL11 and VASH/SVBP (creating αΔTyr, a preferred TTLL11 substrate) is significantly higher compared to TTLL11-only transfected cells. ttTMCP reverses TTLL11-mediated polyglutamylation. TTLL6 and TTLL7 polyglutamylate of α-tubulin and β-tubulin, respectively**. D,E**. **Tubulin polyglutamylation WT and SVBP KO mice brain and cortical neurons** cultured for 8 days *in vitro*. Representative immunoblots and corresponding quantification of the ratio between (polyE α-tubulin)/(total α-tubulin) and (polyE β- tubulin)/(total β-tubulin) in mice brains (**D**), and neurons (**E**). The polyE signal of β-tubulin in neuron lysates was too low to quantify. Data represent mean ± SEM. n = 5 - 6 animals and 6 independent neuronal differentiation experiments of each genotype. Unpaired t test, ns, not significant, ***p < 0.001 and ****p < 0.0001. Comparative analyses of αTyr and αΔTyr variant levels in samples from D and E are shown in Figure S12.

Furthermore, we investigated the impact of TTLL11 ectopic expression in hippocampal neurons cultured *in vitro*. The neurons, which are known to be rich in αΔTyr and αΔ2 variants (*32*) as well as in polyglutamylated tubulins (*33, 34*), demonstrated a loss of αΔ2 following overexpression of TTLL11 (Fig. S12A). These results offer indirect evidence in support of the concept that TTLL11 polyglutamylates the C-terminus of α-tubulin, thereby reversing the αΔ2 to αΔTyr and/or extending the main chain by a C-terminal polyglutamate chain.

Finally, we studied whether the loss of VASH/SVBP enzymes *in vivo* could influence the level of tubulin polyglutamylation in the brain by reducing the availability of αΔTyr, the preferred TTLL11 substrate (Fig. S12B,C). We thus compared the levels of polyglutamylated tubulin in brain extracts and neuronal cultures from WT and SVBP KO mice, given that the defective tubulin detyrosination, resulting from the absence of the chaperone SVBP, has a marked impact on the structure and function of the brain in both humans and mice (*35*). In accordance with the result from biochemical and cellular experiments, the data obtained from *in vivo* studies demonstrated that the absence of VASH/SVBP detyrosinating enzymes markedly diminishes α-tubulin polyglutamylation in both cellular and brain tissue models, whereas the polyglutamylation levels of β-tubulin remain unaltered (Fig. 7D,E).

## Discussion

Together with TTLLs 1, 6, 9, and 13, TTLL11 is classified as glutamate elongase (*19*). The original biochemical findings were later corroborated by comprehensive structural analyses of TTLL6, which identified Gln180 as the primary determinant of its selectivity (*27*). The arrangement of the active site of TTLL11 is analogous to TTLL6, including the conservation of Gln255 corresponding to Gln180 of TTLL6. Our MS/MS data confirmed the classification of TTLL11 as an elongase by showing that elongation is much preferred compared to the lateral branching activity and that elongation is likely the only TTLL11’s physiologically relevant activity. Furthermore, our findings strongly suggest that elongation of tubulin primary chains by TTLL11 is preferred over elongation of preexisting glutamate lateral branches, although the mechanistic basis for this selectivity remains unclear and cannot be derived from the cryo-EM structure.

Earlier TTLL6/7 structures revealed tripartite (or quadripartite) interactions with MTs, involving catalytic domains (yellow), MT-binding domains (MTBD; orange), and MT-binding helices (dark red) (Fig. S4). Moreover, an additional interaction motif must include tubulin C-tails and the residues lining the active sites of TTLL6/7, although the tubulin C-tails are absent from cryo-EM maps likely due to the proposed flexibility and disorder/order transition of the TTLL catalytic domains (*26, 29*). In contrast, the TTLL11/MT interface is much simpler, comprising solely the MT-BHB and the "missing" active site interaction motif. Given the pronounced selectivity of TTLL6 and TTLL7 towards the α- and β-tubulin C-tails, respectively, it seems reasonable to hypothesize that interaction interfaces in TTLL6/7/MT complexes, which are missing in TTLL11, contribute to the protomer selectivity of TTLL6/7.

The sequence homology between TTLL11 and other TTLL elongases is limited to the folded catalytic domain, which terminates at the residue S447 (Fig. S5E). In TTLL6 and TTLL7, the catalytic domain is followed by a helix-loop-helix motif, designated MTBD, and the positively charged helices of MTBD have been shown to interact with MT protomers in the TTLL6/7 cryo-EM structures (*26, 29*). In TTLL11, the corresponding putative MTBD (residues 487 – 526) is absent from the TTLL11 cryo-EM map, suggesting it does not directly interact with the MT surface. The absence of MT interactions is further supported by the overall negatively charged electrostatic surface of the TTLL11 structural motif analogous to MTBDs of TTLL6/7 (Fig. S4). Finally, mutating the cationic patch of the putative MTBD-like motif (the K_488_KKR_491_ to E_488_EEE_491_ mutant), resulted in only a slight reduction in MT binding in our TIRF experiments (Fig. 3B). Taken together, the sequences corresponding to the MTBDs of TTLL6 and 7 do not necessarily constitute the obligatory MT-binding motif in other TTLL elongases and might be dispensable for their physiological functions as shown for TTLL11.

The unique mode of TTLL11 binding to MTs is further illustrated by the superposition of its MT- BHB and corresponding helix pairs of TTLL6/7 (Fig. S4, S6). In TTLL11, helices α11 and α12 of MT-BHB bind the MT protomer groove interface via a mixed ionic/hydrophobic mode, while the positively charged C-terminal end of helix α9 serves as the primary MT-interacting motif in TTLL6 (*29*). These structural differences lead to distinct catalytic domain topologies: TTLL6/7 modify C-tails of tubulin protomers situated toward the plus end of the same protofilament, whereas TTLL11 glutamylates C-tails of protomers positioned toward the minus end of an adjacent protofilament (Fig. 2G, Fig. S4). Taken together, these findings suggest that TTLLs have evolved distinct substrate recognition strategies, albeit with some degree of crosstalk between the catalytic and MT-interacting domains. It would be interesting to explore in future, whether the paradigm of multipartite substrate recognition extends to other TTLLs’ physiological substrates (*20, 22*).

Our structural, biochemical, and functional data provide the following complex picture on TTLL11 preferences for native tubulin C-termini (Fig. 6B). The primary driving force behind the elongation of the tubulin main chain is the peptide sequence and the biophysical characteristics of the C-terminal amino acid. It is evident that (poly)glutamate is preferred, although small aliphatic/hydrophobic residues are also tolerated, whereas bulky residues like tyrosine are excluded (Fig. 5B). Additionally, the protomer type appears to be less critical, with the C-terminal tubulin sequence serving as the primary determinant of TTLL11 substrate specificity.

Importantly, for both α- and β-tubulins, TTLL11 can salvage C-terminally truncated variants that were previously considered "dead-end-sinks". This generates novel polyglutamylation patterns within the tubulin code, directly linking polyglutamylation to the detyrosination/tyrosination cycle. Firstly, as the αΔ2 variant can be reverted to αΔTyr, it can in principle serve as a reservoir for αΔTyr (or αTyr) tubulins in the cell. The spatiotemporal distribution, activity, and crosstalk between all key players of the detyrosination/tyrosination/polyglutamylation pathways are therefore of critical importance in regulating cytoskeleton-dependent cellular processes. For example, it can be assumed that in addition to detyrosination, C-terminal polyglutamylation may serve as yet another mechanism to regulate αTyr levels, thereby modulating processes mediated by αTyr readers, such as CAP-Gly proteins and kinesin-13 depolymerizing motors in the case of intracellular trafficking and MT disassembly, respectively (*36–40*). By polyglutamylating tubulin primary polypeptide chains, TTLL11 emerges as a pivotal player in expanding the landscape of the functional MT-related modifications within the tubulin code.

Polyglutamylation of MTs is essential for regulating physiological processes such as intracellular transport, axonal growth, chromosomal separation, and ciliogenesis (*18, 41–45*). As for the underlying molecular mechanisms, polyglutamylation profoundly impacts the activity of enzymes that modify or interact with MTs. For example, MT-severing enzymes, such as spastin, preferentially bind polyglutamylated MTs (*23*) and polyglutamylation modulates the recruitment and activity of motor proteins like dynein and kinesin. It also affects MT dynamics by controlling the activity of stabilizing or destabilizing MT-associated proteins (MAPs), fine-tuning the structural and functional output of MTs in response to cellular demands (*33, 46–49*). While polyglutamate chain length is known to impact these processes (*50, 51*), it remains to be established whether tubulin code readers are sensitive to the nature of the linkage of glutamate chains, i.e., if they could distinguish between polyglutamylation at branching points vs polyglutamylation extending tubulin primary chains. Overall, data reported here thus provide an impetus for more detailed biochemical and physiological studies focusing on these relationships.

## Materials and Methods

### Experimental Design

#### Statistical analysis

##### Sequences and cloning

Clones of human TTLL11 (hTTLL11; UniProt Q8NHH1), the inactive hTTLL11(E441G) mutant, and murine CCP1 (mCCP1) were a kind gift of prof. Carsten Janke (*19*). The TTL clone was reported previously (*4*). Truncated TTLL11 variants were PCR amplified using corresponding sets of gene-specific primers (Table S1) and cloned into the pDONR221 donor vector *via* the BP recombination reaction following the manufacturer protocol (BP Gateway cloning, Invitrogen). Individual sequences were transferred into the pMM322 expression vector comprising the N-terminal Twin-Strep-FLAG-HALO-tag (*52*) using the LR recombination reaction. Sequence-optimized genes encoding human TCMP2 (Q8NCT3), ttTMCP (*Tetrahymena Thermophila*, Q24D80), human TTLL6 (Q8N841), and human TTLL7 (Q6ZT98) were synthesized commercially (Thermo Fisher Scientific, MA USA) and cloned the pMM322 expression vectors as described above. The TTL gene was PCR amplified using corresponding sets of gene-specific primers (Table S1) and cloned into the pEC566 expression vector containing the N-terminal His-MBP tag (*52*). All used plasmids are listed in Table S1.

Vectors for lentiviral expression (pHR-CMV – expression, psPAX2 – packaging, pMD2.G envelope) were kindly provided by Daniel Rozbesky (*53*). The target genes were cloned into the pHR-CMV expression plasmid using the Gibson Assembly Master Mix (BioLabs Inc., New England) following the manufacturer protocol together with specific primers with overlapping target sequences (Table S1). Nucleotide sequences of all plasmids were verified using Sanger sequencing.

Site-directed mutagenesis was carried out by the standard QuikChange protocol (Agilent) using matching pairs of complementary mutagenic primers (Table S1).

### Protein expression and purification

#### Transduction of HEK293T cells

The published lentiviral transduction system was used to generate stable transformants of HEK293T cells (*53*). Briefly, 10 μg pHR-CMV-TetO2 containing the gene of interest, 10 μg psPAX2, and 10 μg pMD2.G plasmid was mixed in the DMEM/F-12/SFM medium in the total volume of 0.25 mL. 75 μg of linear polyethyleneimine (lPEI; Polysciences Inc., Warrington, PA, USA) in 0.25 mL of DMEM/F-12/SFM was mixed with the plasmid solution and incubated at room temperature for 20 min. The mix was then added to a T75 flask with HEK293T Lenti-X cells at >90 % confluency in 11.5 mL of fresh DMEM/F-12/2 % FBS medium. The cells were incubated at 37 °C, 5% CO2 for 72 hours. The conditioned medium was harvested, mixed with 6 mL of fresh DMEM/F-12/10 % FBS medium and filtered through a 0.45 μm filter unit. HEK293T cells (>90 % confluency) were transduced with the medium containing the lentiviral particles in a T75 flask. The medium was exchanged after 3 days. To enrich positive cell population, the transduced cells were labeled with the 50 nM TAMRA-HALO probe for 30 min, medium replaced and cells sorted using BD Aria Fusion FACS (BD Biosciences, NJ USA). 100 000 cells were collected, transferred to DMEM/F-12/10 % FBS medium and expanded for ensuing experiments.

### Heterologous expression and purification

#### TTLL11 variants

hTTLL11 variants were expressed in suspension HEK293T cells as fusions with the N-terminal TwinStrep-Flag-HALO tag using established protocols (*52, 54*). Briefly, a day prior to transfection, cells were seeded at the density of 2.0 x 10^6^ cells/mL in 350 mL of Free Style F17 medium (Thermo Fisher Scientific) supplemented with 2 mM L-glutamine and 0.1 % Pluronic F-68 (Invitrogen) in 2L Erlenmeyer flasks. The cell suspension was incubated at 110 RPM under a humidified 5 % CO2 atmosphere at 37 °C overnight and the next day transfected with a mixture of 0.7 mg of an expression plasmid DNA diluted in 17.5 mL of PBS and 2.1 mL of lPEI (1mg/mL). 4 hours post-transfection, 350 mL of ExCell serum-free medium (Merck Life Sciences, Darmstadt, Germany) was added to the cell suspension. Cells were harvested by centrifugation at 500 g, 5 mins, 4 °C, 72 hours post-transfection. The cell pellet was lysed in an ice-cold lysis buffer (100 mM Tris-HCl, pH 8.0, 10 mM NaCl, 5 mM KCl, 2 mM MgCl2, 10 % glycerol, 1 mM EDTA, 1 U/mL benzonase (Merck Life Sciences, Darmstadt, Germany), 1x protease inhibitor cocktail (Roche, Basel, Switzerland)) by sonication (24 W/3x20 s min). To assist cell lysis Igepal-630 (0.2 %, v/v) was added to the cell lysate (20-min incubation on ice), followed by the addition of 4M NaCl (150 mM final concentration). The cell lysate was then centrifuged sequentially at 9 000 g (15 mins), 4 °C and 30 000 g (30 mins) at 4 °C.

For purification, the supernatant was mixed with a Strep-Tactin XT resin (IBA, Gottingen, Germany) equilibrated in 100 mM Tris-HCl, pH 8.0, 150 mM NaCl, 5 mM KCl, 2 mM MgCl2, 10 % glycerol, 1 mM EDTA (equilibration buffer) and incubated at 4 °C for 1 hour. The slurry was then loaded into a 10 mL plastic column, and sequentially washed with the ice-cold equilibration buffer (20 column volumes; CV), 5 CVs of the equilibration buffer supplemented with 3 mM ATP and 10 mM MgCl2, and finally 5 CVs of the equilibration buffer. The fusion protein was eluted with 12 CVs the elution buffer (50 mM Tris-HCl, pH 8.0, 150 mM NaCl, 10 mM KCl, 10 % glycerol, 10 mM D-biotin, 1 mM EDTA), and pooled elution fractions concentrated to approximately 2 mg/mL, snap frozen in liquid nitrogen and stored at –80 °C. The sample was centrifuged at 20 000 g, 4 °C, for 15 min, transferred onto a Nanosep 0.2 μm spin filter tube (Pall Corp., Port Washington, NY, USA) and centrifuged at 10 000 g, 4 °C, for 5 min. The supernatant was injected onto a size exclusion Superose 6 Increase 10/300 GL column (GE Healthcare Bio-Sciences Corp., Piscataway, NJ, USA) connected to the NGC Discover Pro System (Bio-Rad Laboratories, Hercules, CA, USA) equilibrated in the SEC buffer (50 mM Tris-HCl, pH 8, 140 mM NaCl, 10 mM KCl, 1mM EDTA, 0.5 mM TCEP, 5 % glycerol). Fractions comprising TTLL11 fusions were pooled, concentrated to approximately 1 mg/mL, snap-frozen in liquid nitrogen and stored at -80°C until further use. The protein purity was evaluated by SDS-PAGE (Fig. S5C) with typical yields ranging from 150 to 800 –μgs per L of cell culture.

### CCP1

Murine CCP1 (mCCP1) and the hTTLL11 (E531G) mutant were expressed in suspension stably transduced HEK293T cells as fusions with the N-terminal TwinStrep-Flag-HALO tag. Cells were grown in the 1:1 mixture of the FreeStyle F17 Expression Medium and the ExCell293 Serum-Free Medium supplemented with 0.05 % Pluronic F-68 and 2 mM L-glutamine at 110 rpm under a humidified 5 % CO2 atmosphere at 37 °C. Once reaching the density of 8.0 x 10^6^ cells/mL, cells were harvested by centrifugation at 500 g at 4°C for 10 mins and recombinant proteins purified by Streptactin affinity chromatography as described above. For CCP1 purification, EDTA was omitted from all purification buffers.

### TTL

TTL was expressed in BL21(DE3) RIPL *E. coli* (Thermo Fisher) as a fusion with the N-terminal His-MBP tag. The cell culture was grown to OD600 = 0.8 at 37 °C, then cooled to 18 °C, and expression was induced by addition of 1 mM IPTG, and cells were cultivated overnight. Cells were harvested by centrifugation (10 000 g, 4 °C, 20 min), and the cell pellet resuspended in buffer A (PBS supplemented with 10 mM MgCl2, 1 mM TCEP), and the protease inhibitor cocktail, pH 7.5. Cells were lysed by three passes through EmulsiFlex (Avestin) reaching 110 bar and then centrifuged subsequently at 7 000 g and 30 000 g, 4 °C, 15 min each step. The fusion protein was purified from the cell supernatant by the Ni-NTA affinity chromatography using a HiTrap™ HP column (Cytiva) connected to FPLC NGCTM Discover 10 Pro. The system was equilibrated in the HT buffer A, the supernatant loaded, and the column washed with the HT buffer A supplemented by 500 mM NaCl, and fractions eluted by a gradient of imidazole (30 mM-500 mM) in 50 mM Tris-HCl, 10 mM MgCl2, 1 mM TCEP, 10 % glycerol, pH = 7.5. Fractions containing the TTL fusion protein were polled, concentrated and diluted in 50 mM Tris-HCl, 10 mM MgCl2, 1 mM TCEP, 10 % glycerol, pH 7.5 to lower the imidazole concentration to 10 mM. The final concentration of TTL was 18 mg/mL, and after flash freezing in liquid nitrogen the protein was stored at -80 °C until further use.

### TTLL11 labeling for TIRF microscopy

The purified HALO-TTLL11 fusions were mixed with a 3-fold molar excess of the Janelia 549-HALO probe (5 mM stock solution in DMSO; Promega) and incubated at 22°C for 30 min. The sample was centrifuged at 20 000 g, 4 °C, for 15 min, transferred onto a Nanosep 0.2 μm spin filter tube (Pall Corp., Port Washington, NY, USA) and centrifuged at 10 000 g, 4 °C, for 5 min. The supernatant was injected onto a size exclusion Superose 6 Increase 10/300 GL column (GE Healthcare Bio-Sciences Corp., Piscataway, NJ, USA) connected to the NGC Discover Pro System (Bio-Rad Laboratories, Hercules, CA, USA) equilibrated in the SEC buffer (50 mM Tris-HCl, 140 mM NaCl, 10 mM KCl, 1mM EDTA, 0.5 mM TCEP, 5 % glycerol, pH 8). Fractions comprising Janelia549-labeled TTLL11 fusions were pooled, concentrated to approximately 1 mg/mL, snap-frozen in liquid nitrogen and stored at -80°C until further use.

### Purification of tubulin from HEK293T cells

Tubulin purification from suspension HEK293T was performed by two polymerization and depolymerization cycles based on a published protocol (*55*). Following the second depolymerization performed in BRB80, the protein was centrifuged, the concentration determined by absorbance measurement at 280 nm using a NanoDrop One Microvolume UV-Vis Spectrophotometer (Thermo Scientific). Purified tubulin was stored in 17 mg/mL aliquots at -80 °C.

### Purification of porcine tubulin

The tubulin purification from pig brain followed a previously published protocol by polymerization/depolymerization cycles similar to tubulin from HEK293T cells (*56*). The concentration was determined by absorbance measurement at 280 nm using a NanoDrop One Microvolume UV-Vis Spectrophotometer (Thermo Scientific). The purified tubulin (20mg/mL) was stored in aliquots in BRB80 buffer at -80 °C.

### Preparation of α-tubulin variants

#### Tyrosinated tubulin

For tyrosination of tubulin isolated from HEK293T cells, a reaction mixture was prepared containing 50 μM tubulin, 4 μM TTL, 300 μM tyrosine, and 2 mM ATP in BRB80 (80 mM PIPES, pH 6.9, 1mM EGTA, 1mM MgCl2). The reaction was incubated at 22°C for 2 hours, followed by centrifugation at 20 000 g for 10 minutes at RT. The resulting supernatant was aliquoted, snap-frozen in liquid nitrogen, and stored at - 80 °C until further use.

#### αΔTyr, αΔ2 and αΔ3 tubulins

To prepare detyrosinated and Δ3 tubulin, a reaction mixture comprising 50 μM tubulin from HEK293T cells, 4 μM carboxypeptidase A (CPA, Merck Life Sciences, Darmstadt, Germany), and 3 μM recombinant mCCP1 (only in Δ3 tubulin preparations) in BRB30 supplemented with 1 μM ZnCl2 and 4 mM MgCl2 was incubated at 35 °C for 1 hour. The reaction mixtures were then supplemented with additional spike of equal dose of CPA (and mCCP1) and incubated for an additional 2 hours. To prepare polymerized MTs and to remove mCCP1, reaction mixtures were supplemented with PIPES-Na at final concentration of 330 mM (1 M stock solution, pH 6.9), and 1 mM GTP, and incubated at 37 °C for 30 minutes. Polymerized MTs were centrifuged at 14 000 g, 35 °C, for 30 minutes, resuspended in BRB80, aliquoted, snap-frozen in liquid nitrogen, and stored at -80 °C until further use.

#### Double stabilized microtubules

Tubulin variants were polymerized as described previously (*57*). Briefly, tubulin aliquots stored at -80 °C were thawed and centrifuged at 21 000 g, 4 °C, for 10 min. The supernatant was transferred to a fresh Eppendorf tube, diluted (to 2.5-10 μM concentration) with the polymerization buffer (BRB80, 1 mM GMPCPP (Jena Bioscience, Jena, Germany), 1 mM MgCl2), incubated on ice for 5 min and then at 37°C for minimum 1 hour. The sample was centrifuged at 14 000 g, 25°C, for 30 min, the supernatant carefully discarded and pelleted MTs dissolved in the BRB80 buffer supplemented with 10 μM Taxol to the final concentration of 10 μM. The double-stabilized MTs (dsMTs) were stored at room temperature until further use.

#### Tubulin glutamylation assay *in vitro*

Glutamylation reactions comprised 1 μM dsMTs and 0.3 μM TTLL11 variants in the BRB30 buffer supplemented with 1 mM TCEP, 2 mM ATP, 15 mM glutamate (L-glutamic acid monosodium salt monohydrate, Merck, Germany; L-glutamic acid 2,3,3,4,4-D5, 98 %, Cambridge Isotope Laboratories, USA; or ^18^O-labelled glutamic acid synthesized in house), 4 mM MgCl2, and 0.5 mM EDTA. The mixture was incubated in an orbital shaker (300 rpm; Thermomixer Comfort, Eppendorf, Germany) at 25°C, for a defined time in the range of 10 min – 2 hours. For the SDS-PAGE/Western blotting analyses, the reaction was stopped by the addition of the 1/5 volume of 5x SDS-PAGE sample buffer (250 mM Tris-HCl, pH 6.8, 10 % w/v SDS, 30 % v/v glycerol, 5 % v/v 2-mercaptoethanol, 0.02 % w/v bromphenol blue) supplemented with 250 mM DTT. For mass spectrometry analyses, the reaction was centrifuged as described below.

#### Tubulin glutamylation in cells

HEK293T cells were transfected with expression plasmids using the transfection reagent JetOptimus (Polyplus, #101000025) according to the manufacturer’s instructions. Cells were lysed 24 hours after transfection by sonication in PBS. 1x SDS-PAGE sample buffer and 200 mM DTT was added to the samples and the samples were incubated at 60°C for 10 minutes followed heating at 95 °C for 5 min. The samples were analyzed by SDS-PAGE and Western blotting.

#### SDS-PAGE, Western blotting

α- and β- tubulins were resolved by SDS-PAGE composed of a 10 % acrylamide separation gel (375 mM Tris-HCl, pH 9, 0.1 % SDS (Sigma #L5750)) and 3.5 % stacking gel (125 mM Tris-HCl, pH 6.8, 0.1 % SDS) prepared using the acrylamide/bis-acrylamide stock solution [40 % acrylamide solution (Bio-Rad #161-0140) mixed with 0.54 % w/v bis-acryl amide dry powder (Bio-Rad #161-0210)]. The running buffer was composed of 50 mM Tris-HCl, 384 mM glycine, and 0.1 % SDS (*58*). The separation was performed at 150 V for 80 min.

For immunoblotting, the proteins were transferred onto a PVDF membrane using the Bio-Rad Trans-Blot Turbo Transfer System (Hercules, CA) under standard conditions. The membrane was blocked with 5 % w/v skimmed milk dissolved in TBS. The membranes were probed with antibody solutions in 2.5 % w/v skimmed milk in TBS (dilutions detailed in Table S1), at 4 °C for 8 h and 2h for the primary and secondary Abs, respectively. Secondary antibodies conjugated with horse radish peroxidase (HRP; Bio-Rad Laboratories) were used for the detection of chemiluminescence signals (Immobiolon Forte Western HRP Substrate, Millipore, MA) and visualized using the ImageQuant LAS 4000 (GE healthcare). The PageRuler Plus Prestained Ladder (Thermo Fisher) was used as a molecular weight marker.

#### Mass spectrometry analysis of glutamylation

dsMTs from glutamylation assays were centrifuged at 14 000 g, 22°C, for 30 min. The resulting pellet, containing 26 μg of MTs, was resuspended in 5 μL 50 mM ammonium bicarbonate, pH 7.8, supplemented with 100 mM DTT. Following incubation at 60 °C for 30 min, freshly prepared 50 mM iodoacetamide (300 mM stock solution; 0.7 µL) was added and the mixture incubated in the dark for 30 min. Next, 150 mM DTT (1 M stock solution; 0.8 µL) and 0.65 μg (AspN; 2.5 µL) or 1 μg (Trypsin/LysC mix; 1 µL) were added and incubated at 37°C (shaking at 650 rpm) overnight.

HPLC separation was performed using an Agilent 1290 series HPLC system (Agilent Technologies, Santa Clara, CA). The sample (5 μL) was injected onto a reverse-phase trap column (Luna Omega Polar C18, 0.3 × 30 mm, Phenomenex, Torrance, CA) followed by a reverse-phase analytical column (Luna Omega Polar C18, 0.3 × 150 mm, Phenomenex), both heated to 50 °C. Mobile phases used: A (2 % acetonitrile, ACN; 0.1 % formic acid (FA)) and B (98 % ACN, 0.1 % FA), the flow rate 10 μL/min. The LC run consisted of a 35 min separation gradient of 5–40 % B, a 3 min spike of 40–95 % B, 3 min washing (95 % B), a 1 min drop of 95–2 % of B, and equilibration of columns in 2 % B for 10 min.

MS (quantification) and MS/MS (fragmentation) analyses were performed using a trapped ion mobility-quadrupole time-of-flight mass spectrometer (timsTOF Pro, Bruker Daltonics, Billerica, MA). Eluted peptides were analyzed by an MS acquisition method recording spectra from 250 to 2500 m/z, and ion mobility was scanned from 0.6 to 1.73 Vs/cm^2^. For MS/MS, the protocol included a TIMS survey scan of 100 ms followed by ten PASEF MS/MS scans, 150 ms for each of ion accumulation and ramp time. The total cycle time was 1.16 s. Target intensity was 20 000, the intensity threshold was 2500, and singly charged peptides with m/z < 800 were excluded by an inclusion/exclusion polygon filter applied within the ion mobility m/z heatmap. Precursors for data-dependent acquisition were fragmented with an ion mobility-dependent collision energy, which was linearly increased from 20 to 59 eV.

Intensities of peptides were determined using DataAnalysis 5.0 software (Bruker Daltonics). Visualization was performed in GraphPad Prism 8 (GraphPad Software, San Diego, CA). Experimental data points represent mean values ± S.D.; *n* = 3. Fragmentation spectra were also analyzed in the same software, fragment masses were determined using GPMAW 12.20 (General Protein/Mass Analysis for Windows) with addition of the mass values of present isotopes (^13^C, D, ^18^O). Masses of the labelled peptides selected for fragmentation and peptide envelope shapes were compared with theoretical spectra at https://envipat.eawag.ch. Due to the length of some analyzed peptides, fragments containing one or two ^13^C were used for the analysis in some cases. Fragments from b and y series were identified showing the precise location of polyglutamate chains in each peptide.

#### TIRF microscopy

Total internal reflection fluorescence (TIRF) microscopy in a combination with an interference reflection microscopy (*59*) was performed using an inverted microscope (Nikon-Ti E, Nikon-Ti2 E) (IRM, Nikon TI2-D-LHLED), equipped with a 100x NA 1.49 oil immersion objective (SR Apo TIRF, Nikon) and a PRIME BSI camera (Teledyne Photometrics). Porcine dsMTs were visualized using the IRM, while Janelia 549 labeled TTLL11 variants were visualized with a 561 nm laser. Exposure time was 50 ms with the laser power set to 10 %. The microscope was controlled using Nikon NIS Elements software. All experiments were conducted at 22°C. Image analysis was carried out in the Fiji software(*60*).

Flow cells were prepared as described previously (*52, 61*). Briefly, microscope chambers were built from silanized coverslips (Corning Cover Glass Product) prepared as described previously (*62*). Parafilm was used to space two glasses and to form channels of ∼0.1-mm thickness, 3-mm width, and 18-mm length. MTs were attached to the glass surface in each chamber *via* an anti-β-tubulin antibody (Sigma-Aldrich, T7816, 20 μg/mL in PBS). Subsequently, Janelia 549 TTLL11 variants were added to the chamber in the binding buffer (40 mM Tris-HCl, pH 7.0, 1mM TCEP, 1mM MgCl2, 5 % v/v glycerol), 0.5 mg/mL casein, 10 μM paclitaxel, 0.001 % C12E8 (dodecyloctaglycol), 20 mM D-glucose, 110 μg/mL glucose oxidase, and 20 μg/mL catalase. The fluorescent signal of Janelia 549-labeled TTLL11 variants was co-localized with the MTs viewed in the IRM channel. The signal intensity was quantified in Fiji by creating a linear ROI with 3 pixels width along the MT. Then the background with the same area near the MT was subtracted and intensities per µm of MT were calculated. Intensities were normalized to the mean signal of the control and visualized using GraphPad Prism.

### Cryo-EM of the TTLL/MT complex

#### Sample preparation

dsMTs were centrifuged at 14 000 g, 25°C, for 30 min and resuspended to 50 μM concentration in the EM buffer (40 mM Tris-HCl, pH 7.0, 1 mM TCEP, 1 mM MgCl2, 5 % glycerol). The final mixture comprised 10 μM TTLL11 (E441G) and 10 μM dsMTs in the EM-buffer. Cryo-EM grids Quantifoil^TM^ R 2/1, Cu 300 mesh (Quantifoil, Germany) were glow discharged using a Leica EM ACE600 apparatus (Leica Microsystems, Germany) for 30 s. A Leica EM GP2 Automatic Plunge Freezer (Leica, Germany) was used with the settings: 30 sec wait time, 4 sec blotting time, and automatic detection of the blotting force. Grids were first scanned using a JEOL JEM 2100-Plus 200kV microscope (JEOL, Germany) and the best grids then used for data collection.

#### Data collection and processing

8.8k movies were collected using Titan Krios (G1-2) - 300kV (Thermo Fisher) equipped with a Falcon 3EC direct electron camera (Thermo Fisher). The data was acquired with the following setup: kV=300, cs=2.7, apix=1.349, frameDose=1.0 e/A^2^ (30 frames), nominal defocus range -0.8 through -2.0 μm (Fig. S3A,B).

Data processing was carried out using the Microtubule Relion-based pipeline (MiRP) (*63*). The 14- protofilament fraction of the data was using the MiRP templates and assigned positions of homogenous segments by a series of MiRP 3D classifications. Initial refinements were performed in RELION using a template map from prior 3D classification, the optimized reference map from the refinement, and finally a helical symmetrization. As evaluating different refinements protocols for the whole MT map did not result in map improvements, symmetry expansion followed by particle subtraction was carried out for smaller segments of the map (from the 12 tubulin units down to an isolated TTLL11 molecule) to account for the presumed flexibility of TTLL11 at the MT surface. Different classifications were tested for each set of particles and the refinement using multiple masks of different sizes and edge expansions was carried out both in Relion (*64*) and Cryosparc (*65*). Final data processing steps were performed in Cryosparc (*65*) and final refinements were done using series of local searches exploiting masks containing either twelve or four decorated tubulin units. For post-processing, we used DeepEMhancer with the highRes setting (*66*). Fourier Shell Correlation (FSC) curves in Cryosparc were used to estimate resolutions of each reconstruction using the gold standard. To estimate resolutions for different parts of the map, a Local Resolution job ran in Cryosparc. The tubulin part of the map has a maximum resolution of 2.7 Å, while the TTLL11 molecule is covered by a resolution gradient from 3.3 to 7 Å (Fig. 2C).

### Model Building

Initial atomic models were prepared in Coot adapted for cryo-EM (*67*) by mutating residues of existing GMPCPP-bound tubulin models (PDB ID 6E7B) to match the human TUBB-5 and TUBA-1B sequences. For TTLL11, an AlphaFold model of human TTLL11 was used. For atomic model building and refinement, a tubulin tetramer and TTLL11 were first docked into the density maps using rigid body fitting in ChimeraX (*68*). The structures were further refined using real space refinement the Phenix suite (*69*) interspersed with the manual corrections to the model in Coot. The quality of the models was validated using the MolProbity server (*70*) and the model deposited in the RCSB under the accession number 9HQ4. Complete model statistics are reported in Table S2.

### Assays on SVBP knock-out mice brain and neurons

#### Animals

All experiments were conducted in accordance with the policy of the Grenoble Institute des Neurosciences (GIN). In compliance with the French legislation and European Union Directive of 22 September 2010 (2010/63/UE), the research was authorized by the Direction Départementale de la protection des populations—Prefecture de l’Isère-France and the ethics committee of GIN n° 004 accredited by the French Ministry of Research. SVBP-deleted mice (C57BL/6) were genotyped by PCR amplification(*35*). Primers for testing SVBP mouse strain were 5′-GATCCACCTGCCCGGAAA-3′, 5′- TTTCTTCCAGCACCCTCTCC-3′ and 5′-CAAACCATGGATCCACGAAA-3′ as already described(*36*).

### Brain protein extracts preparation

Fifteen-week-old mice were sacrificed by cervical dislocation. Cerebral hemispheres were extracted, rinsed in phosphate buffered saline solution, shock frozen in liquid nitrogen and stored in liquid nitrogen until use. The organs (∼200 mg) were lysed in 1 ml PEM buffer (100 mM PIPES, 1 mM EGTA, 1 mM MgCl2 and protease inhibitor cocktail (complete mini^TM^, Roche Diagnostics) at pH 6.7) using a Bead Mill 4 Fischer brand system with 1,4 mm ceramic beads. The lysate was centrifuged at 17 000 g for 30 min at 4 °C. The pellet was homogenized in 1 ml SDS-buffer (3% SDS, 30 mM Tris-base, pH 8.8, 5 mM EDTA, 30 mM NaF, 10 % glycerol and 1 mM DTT) and centrifuged at 17000 g for 15 min at 20°C. The supernatant was used for immunoblotting assays.

### Neuronal cell culture and transfection

Mouse hippocampi (immunofluorescence) or cortices (immunoblot) were dissected from embryos (E17.5) and digested in 0.25% trypsin in Hanks’ balanced salt solution (HBSS, Invitrogen, France) at 37°C for 15 minutes. After manual dissociation, neurons were plated on 0.5-1 mg/ml poly-L- lysine-coated dishes, incubated 2 h in DMEM-10% horse serum and then changed to magnetic cell sorting (MACS) neuro medium (MiltenylBiotec) with B27 supplement (Invitrogen, 486 France). Neurons were transfected using Amaxa Nucleofector kits (Lonza). At day 0 of culture, hippocampal cells from one or two embryos were used for a transfection point with 3 µg of TTLL11-EYFP DNA (kind gift of Janke lab, described in Van Dijk et al (*19*) and then cultured for 2 or 4 days. For immunoblotting, cortical cells cultured 8 days *in vitro* were directly collected in the SDS-PAGE sample buffer.

### Antibodies, immunoblotting and immunofluorescence

Antibodies specific for total tubulin (mouse, α3A1 or β3A11), tyrosinated-tubulin (rat, YL1/2), detyrosinated-tubulin (rabbit) and Δ2-tubulin (rabbit) were described by Aillaud et al (*12*), chicken anti-GFP antibody was purchased from Sigma. A novel polyclonal antibody against poly-glutamates chain (anti-polyE Gre) was produced in guinea pig by using the peptide C-EEEEEEE linked at the N-terminal to the keyhole limpet hemocyanin protein via the cysteine. Validation of this antibody was performed in protein extracts from HEK293T cells overexpressing TTLL enzyme. For immunoblotting, antibodies were all used at 1:10000, except anti-polyE Gre which was used at 1:5000. For immunofluorescence, antibodies were all used at 1:1000, except α3A1 which was used at 1:5000.

For immunoblotting, protein extracts were loaded on 10% acrylamide gels (Mini-PROTEAN® TGX Stain- Free™, Biorad) or home-made 10% acrylamide/bisacrylamide (37.5 :1) for separation of α- and β-tubulin, and transferred with Trans-Blot® Turbo (Bio-Rad) on PVDF membranes. Membranes were incubated with primary antibodies, with secondary fluorescent antibodies (anti mouse conjugated to Alexa-488, and anti-guinea pig, anti-rabbit, and anti-rat conjugated to Cyanine 3, all used at 1:2000) and finally revealed with Chemidoc camera (Bio-Rad). The immunoreactive bands were visualized by ChemiDoc MP Imaging System (Bio-Rad), and then images were quantified and analyzed using Image Lab (Bio-Rad) software. The quantifications were graphically plotted using GraphPad software.

For immunofluorescence, cells were fixed at 37°C in 4% paraformaldehyde, 4.2% sucrose, phosphate buffered saline medium (PBS) and permeabilized using 0.1% Triton X-100, PBS. Cells were then incubated with primary antibodies, followed by incubation with anti-chicken conjugated to Alexa-488, anti-mouse conjugated to Alexa-647 or anti-rabbit conjugated Cyanine-3 (all at 1:500). Nuclei were stained using Hoescht 33258 (1 μg/ml).

#### Synthesis of ^18^O-labeled glutamate

All solvents used for synthesis were obtained from commercial sources. All chemicals were purchased from Sigma-Aldrich, TCI, Combi-Blocks or Armar isotopes (water-^18^O) and were used without further purification. TLC was performed on Silica gel 60 F254-coated aluminum sheets (Merck). Products were purified by preparative scale HPLC on a JASCO PU-975 instrument (flow rate 10 ml/min) equipped with a UV-975 UV detector and Waters YMCPACK ODS-AM C18 Prep Column (5 µm, 20 × 250 mm). The purity of compounds was assessed on an analytical JASCO PU-1580 HPLC (flow rate 1 ml/min, invariable gradient from 2 to 100 % acetonitrile in 30 min) with a Watrex C18 Analytical Column (5 µm, 250 × 5 mm). ^1^H and ^13^C NMR spectra were measured using Bruker AVANCE III HD 400 MHz, Bruker AVANCE III HD 500 MHz, and Bruker AVANCE III 600 MHz instruments. The internal signal of TMS (δ 0.0, CDCl3), residual signal of CDCl3 (δ 7.26), or D2O (δ 4.79) were used for standardization of ^1^H NMR spectra. NMR spectra were recorded at room temperature unless noted otherwise. Chemical shifts are given in δ scale; coupling constants (*J*) are given in Hz. The ESI mass spectra were recorded using a ZQ micromass mass spectrometer (Waters) equipped with an ESCi multimode ion source and controlled by MassLynx software. Low-resolution ESI mass spectra were recorded using a quadrupole orthogonal acceleration time-of-flight tandem mass spectrometer (Q-Tof micro, Waters) and high-resolution ESI mass spectra using a hybrid FT mass spectrometer combining a linear ion trap MS and Orbitrap mass analyzer (LTQ Orbitrap XL, Thermo Fisher Scientific). The conditions were optimized for suitable ionization in the ESI Orbitrap source (sheath gas flow rate of 35 au, aux gas flow rate of 10 au of nitrogen, source voltage of 4.3 kV, capillary voltage of 40 V, capillary temperature of 275 °C, tube lens voltage of 155 V). The samples were dissolved in methanol and applied by direct injection.

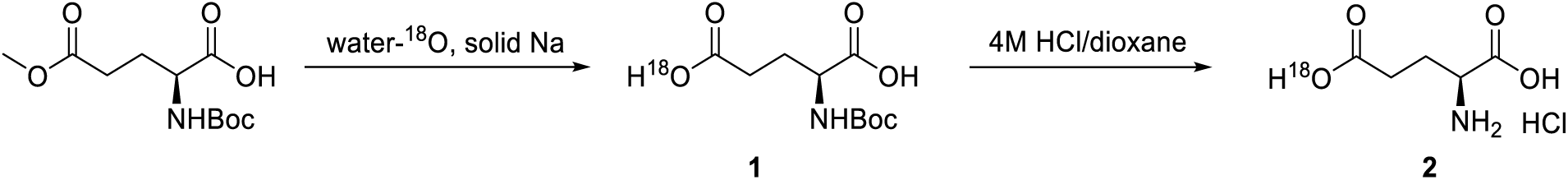

### (tert-Butoxycarbonyl)-L-glutamic-γ-^18^O acid (1)

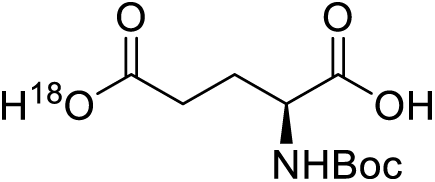

Sodium metal (11.0 mg, 0.48 mmol, 3.0 equiv.) was carefully dissolved in water- ^18^O (0.1 mL) under argon atmosphere at 0° C. This solution was added to (*S*)-2- ((*tert*-butoxycarbonyl)amino)-5-methoxy-5-oxopentanoic acid (42 mg, 0.16 mmol, 1.0 equiv.) and the resulting mixture was stirred under argon atmosphere at room temperature for 1 hour. Reaction mixture was then freezed and lyophilized to afford 40 mg (quantitative yield) of (*tert*-butoxycarbonyl)-L-glutamic-γ-^18^O acid (**1**) as white solid. This intermediate was used for the next step without further purification.

**ESI MS**: 272.1 ([M + Na]^+^).

### L-glutamic-γ-^18^O acid hydrochloride (2)

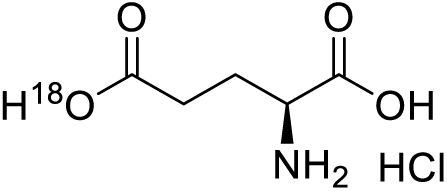

(*tert*-Butoxycarbonyl)-L-glutamic-γ-^18^O acid (**1**, 40 mg, 0.16 mmol) was dissolved in 4M HCl/dioxane (0.2 mL) and the resulting mixture was stirred under argon atmosphere at room temperature for 2 hours. Reaction mixture was then frozen and lyophilized to afford 29 mg (quantitative yield) of L-glutamic- γ-^18^O acid hydrochloride (**2**) as white solid.

^1^H NMR (401 MHz, D2O): δ 2.10 – 2.25 (m, 2H), 2.59 (td, *J* = 2.7, 7.8 Hz, 2H), 3.89 (t, *J* = 6.5 Hz, 1H). ESI MS: 148.0 ([M - H]^+^).

HR ESI MS: calculated for C5H8O3^18^ON: 148.05013; found: 148.04988.

## Supporting information

Supplement Materials Descrition

## Acknowledgments

We thank B. Havlinova, I. Jelinkova, J-M. Soleilhac, and zootechnicians of the Grenoble Institute of Neurosciences for animal care for their technical assistance; P. Pompach for MS analyses; J. Miksatko, J. Novacek, and L. Gaifas for cryo-EM technical support; S. R. Rios and A. Zorgniotti for help with neuronal cultures and immunoblotting; G. Schoehn and A. Peuch for maintaining the IBS/ISBG EM platform and the EM computing cluster.

## Funding

Czech Academy of Sciences (RVO: 86652036)

Czech Science Foundation (GA23-07149S)

Grant agency of Charles University (Projects Nos: 1414120, 284023)

International mobility grant number (CZ.02.2.69/0.0/0.0/18_053/0016973)

Instruct-ERIC - centre FR2, ISBG-IBS (PID 1726)

CF CryoEM of CIISB, Instruct-CZ Centre, and European Regional Development Fund-Projects „Innovation of CIISB“ and „UP CIISB“ (No. CZ.02.01.01/00/23_015/0008175, No. CZ.02.1.01/0.0/0.0/18_046/0015974) supported by MEYS CR (LM2023042)

IMCF at BIOCEV supported by MEYS CR (LM2023050 Czech-BioImaging)

IBS Grenoble EM facility (the Grenoble Instruct-ERIC centre, ISBG; UAR 3518 CNRS-CEA-UGA-EMBL) within the Grenoble Partnership for Structural Biology, supported by FRISBI (ANR-10-INBS-0005-02) and GRAL, financed within the University Grenoble Alpes graduate school (Ecoles Universitaires de Recherche) CBH-EUR-GS (ANR-17-EURE-0003). The EM facility is supported by the Rhône-Alpes Region, the Fondation Recherche Medicale (FRM), the fonds FEDER and the GIS-Infrastrutures en Biologie Sante et Agronomie (IBISA).

Leducq Foundation (grant n° 20CVD01 to M.-J. Moutin) by Institut National de la Santé et de la Recherche Médicale (INSERM), Centre National de la Recherche Scientifique (CNRS), University Grenoble Alpes

Agence National de la Recherche (grant SPEED-Y n° ANR-20-CE16-0021 to M.-J. Moutin)

Photonic Imaging Center of Grenoble Institute Neuroscience (Univ Grenoble Alpes – Inserm U1216), which is part of the ISdV core *facility*

### Author contributions

Conceptualization and experiment design: CB, JC

TIRF Microscopy: JC, MaV, LM, KU

Protein purification: JC, MiV CryoEM: JC, AD, IG

Model building and structure analysis: JC, CB

Mass spectrometry: JC, JK

Biochemical experiments: JC, MiV, MB, ZN

Cell-based glutamylation asseys: MaV

Mice and neuronal culture experiments: SI, MJM

Writing – original draft: CB, JC

Writing – review and editing: CB, JC, IG, MJM, AD, ZN, MaV, LM

## Declaration of generative AI and AI-assisted technologies in the writing process

During the preparation of this work, the authors used ChatGPT in order to ensure good comprehension of the text.

## References

1. A. Roll-Mecak, How cells exploit tubulin diversity to build functional cellular microtubule mosaics. Curr Opin Cell Biol 56, 102–108 (2019).

2. A. Roll-Mecak, The Tubulin Code in Microtubule Dynamics and Information Encoding. Dev Cell 54, 7–20 (2020).

3. C. i. https://BioRender.com, BioRender: Scientific Image and Illustration Software. (2024).

4. C. Aillaud et al., Vasohibins/SVBP are tubulin carboxypeptidases (TCPs) that regulate neuron differentiation. Science 358, 1448–1453 (2017).

5. J. Nieuwenhuis et al., Vasohibins encode tubulin detyrosinating activity. Science 358, 1453–1456 (2017).

6. L. Landskron et al., Posttranslational modification of microtubules by the MATCAP detyrosinase. Science 376, (2022).

7. S. Nicot et al., A family of carboxypeptidases catalyzing α- and β-tubulin tail processing and deglutamylation. Sci Adv 9, (2023).

8. K. Rogowski et al., A Family of Protein-Deglutamylating Enzymes Associated with Neurodegeneration. Cell 143, 564–578 (2010).

9. O. Tort et al., The cytosolic carboxypeptidases CCP2 and CCP3 catalyze posttranslational removal of acidic amino acids. Mol Biol Cell 25, 3017–3027 (2014).

10. D. Raybin, M. Flavin, An enzyme tyrosylating alpha-tubulin and its role in microtubule assembly. Biochem Biophys Res Commun 65, 1088–1095 (1975).

11. L. Paturle-Lafanechere et al., Accumulation of delta 2-tubulin, a major tubulin variant that cannot be tyrosinated, in neuronal tissues and in stable microtubule assemblies. J Cell Sci 107 **( Pt** **6**), 1529–1543 (1994).

12. C. Aillaud et al., Evidence for new C-terminally truncated variants of α- and β-tubulins. Molecular Biology of the Cell 27, 640–653 (2016).

13. M. M. Magiera, P. Singh, S. Gadadhar, C. Janke, Tubulin Posttranslational Modifications and Emerging Links to Human Disease. Cell 173, 1323–1327 (2018).

14. J. E. Lee et al., CEP41 is mutated in Joubert syndrome and is required for tubulin glutamylation at the cilium. Nat Genet 44, 193–199 (2012).

15. P. Xia et al., Glutamylation of the DNA sensor cGAS regulates its binding and synthase activity in antiviral immunity. Nat Immunol 17, 369–378 (2016).

16. A. Deshpande et al., TTLL12 has a potential oncogenic activity, suppression of ligation of nitrotyrosine to the C-terminus of detyrosinated alpha-tubulin, that can be overcome by molecules identified by screening a compound library. PLoS One 19, e0296960 (2024).

17. L. Froidevaux-Klipfel et al., Septin cooperation with tubulin polyglutamylation contributes to cancer cell adaptation to taxanes. Oncotarget 6, 36063–36080 (2015).

18. I. Zadra et al., Chromosome segregation fidelity requires microtubule polyglutamylation by the cancer downregulated enzyme TTLL11. Nature Communications 13, (2022).

19. J. van Dijk et al., A targeted multienzyme mechanism for selective microtubule polyglutamylation. Mol Cell 26, 437–448 (2007).

20. M. Kravec, et al., A new mechanism of posttranslational polyglutamylation regulates phase separation and signaling of the Wnt pathway protein Dishevelled. (2023).

21. B. M. Lorton et al., Glutamylation of Npm2 and Nap1 acidic disordered regions increases DNA mimicry and histone chaperone efficiency. iScience 27, 109458 (2024).

22. J. van Dijk et al., Polyglutamylation is a post-translational modification with a broad range of substrates. J Biol Chem 283, 3915–3922 (2008).

23. B. Lacroix et al., Tubulin polyglutamylation stimulates spastin-mediated microtubule severing. J Cell Biol 189, 945–954 (2010).

24. R. O’Hagan et al., Glutamylation Regulates Transport, Specializes Function, and Sculpts the Structure of Cilia. Curr Biol 27, 3430–3441 e3436 (2017).

25. H. Mathieu et al., Genetic variant of TTLL11 gene and subsequent ciliary defects are associated with idiopathic scoliosis in a 5-generation UK family. Sci Rep 11, 11026 (2021).

26. Christopher P. Garnham et al., Multivalent Microtubule Recognition by Tubulin Tyrosine Ligase-like Family Glutamylases. Cell 161, 1112–1123 (2015).

27. K. K. Mahalingan et al., Structural basis for polyglutamate chain initiation and elongation by TTLL family enzymes. Nat Struct Mol Biol 27, 802–813 (2020).

28. G. Fu et al., Integrated regulation of tubulin tyrosination and microtubule stability by human alpha-tubulin isotypes. Cell Rep 42, 112653 (2023).

29. K. K. Mahalingan et al., Structural basis for alpha-tubulin-specific and modification state-dependent glutamylation. Nat Chem Biol, (2024).

30. L. A. Amos, Microtubule structure and its stabilisation. Org Biomol Chem 2, 2153–2160 (2004).

31. . . (Sep 30th 2024), vol. 2024.

32. C. Sanyal et al., The detyrosination/re-tyrosination cycle of tubulin and its role and dysfunction in neurons and cardiomyocytes. Semin Cell Dev Biol 137, 46–62 (2023).

33. C. Janke, M. M. Magiera, The tubulin code and its role in controlling microtubule properties and functions. Nat Rev Mol Cell Biol 21, 307–326 (2020).

34. K. Natarajan, S. Gadadhar, J. Souphron, M. M. Magiera, C. Janke, Molecular interactions between tubulin tails and glutamylases reveal determinants of glutamylation patterns. EMBO reports 18, 1013–1026 (2017).

35. A. T. Pagnamenta et al., Defective tubulin detyrosination causes structural brain abnormalities with cognitive deficiency in humans and mice. Hum Mol Genet 28, 3391–3405 (2019).

36. A. Konietzny et al., Efficient axonal transport of endolysosomes relies on the balanced ratio of microtubule tyrosination and detyrosination. J Cell Sci 137, (2024).

37. J. J. Nirschl, M. M. Magiera, J. E. Lazarus, C. Janke, E. L. Holzbaur, alpha-Tubulin Tyrosination and CLIP-170 Phosphorylation Regulate the Initiation of Dynein-Driven Transport in Neurons. Cell Rep 14, 2637–2652 (2016).

38. L. Peris et al., Tubulin tyrosination is a major factor affecting the recruitment of CAP-Gly proteins at microtubule plus ends. J Cell Biol 174, 839–849 (2006).

39. L. Peris et al., Motor-dependent microtubule disassembly driven by tubulin tyrosination. J Cell Biol 185, 1159–1166 (2009).

40. M. O. Steinmetz, A. Akhmanova, Capturing protein tails by CAP-Gly domains. Trends Biochem Sci 33, 535–545 (2008).

41. S. Bodakuntla et al., Tubulin polyglutamylation is a general traffic-control mechanism in hippocampal neurons. J Cell Sci 133, (2020).

42. Y. M. Lu, S. Yan, S. C. Ti, C. Zheng, Editing of endogenous tubulins reveals varying effects of tubulin posttranslational modifications on axonal growth and regeneration. Elife 13, (2024).

43. N. Pathak, T. Obara, S. Mangos, Y. Liu, I. A. Drummond, The zebrafish fleer gene encodes an essential regulator of cilia tubulin polyglutamylation. Mol Biol Cell 18, 4353–4364 (2007).

44. A. M. Sheikh, S. Tabassum, Potential role of tubulin glutamylation in neurodegenerative diseases. Neural Regen Res 19, 1191–1192 (2024).

45. D. Ten Martin et al., Tubulin glutamylation regulates axon guidance via the selective tuning of microtubule-severing enzymes. EMBO J, (2024).

46. S. Chakraborti, K. Natarajan, J. Curiel, C. Janke, J. Liu, The emerging role of the tubulin code: From the tubulin molecule to neuronal function and disease. Cytoskeleton (Hoboken) 73, 521–550 (2016).

47. S. Gadadhar, S. Bodakuntla, K. Natarajan, C. Janke, The tubulin code at a glance. Journal of Cell Science, (2017).

48. E. D. McKenna, S. L. Sarbanes, S. W. Cummings, A. Roll-Mecak, The Tubulin Code, from Molecules to Health and Disease. Annu Rev Cell Dev Biol 39, 331–361 (2023).

49. G. A. Viar, G. Pigino, Tubulin posttranslational modifications through the lens of new technologies. Curr Opin Cell Biol 88, 102362 (2024).

50. M. Sirajuddin, L. M. Rice, R. D. Vale, Regulation of microtubule motors by tubulin isotypes and post-translational modifications. Nat Cell Biol 16, 335–344 (2014).

51. M. L. Valenstein, A. Roll-Mecak, Graded Control of Microtubule Severing by Tubulin Glutamylation. Cell 164, 911–921 (2016).

52. L. Skultetyova et al., Human histone deacetylase 6 shows strong preference for tubulin dimers over assembled microtubules. Sci Rep 7, 11547 (2017).

53. J. Elegheert et al., Lentiviral transduction of mammalian cells for fast, scalable and high-level production of soluble and membrane proteins. Nat Protoc 13, 2991–3017 (2018).

54. Z. Kutil et al., The unraveling of substrate specificity of histone deacetylase 6 domains using acetylome peptide microarrays and peptide libraries. FASEB J 33, 4035–4045 (2019).

55. J. Souphron et al., Purification of tubulin with controlled post-translational modifications by polymerization-depolymerization cycles. Nat Protoc 14, 1634–1660 (2019).

56. M. Castoldi, A. V. Popov, Purification of brain tubulin through two cycles of polymerization-depolymerization in a high-molarity buffer. Protein Expr Purif 32, 83–88 (2003).

57. K. Ustinova et al., The disordered N-terminus of HDAC6 is a microtubule-binding domain critical for efficient tubulin deacetylation. J Biol Chem 295, 2614–2628 (2020).

58. M. M. Magiera, C. Janke, Investigating tubulin posttranslational modifications with specific antibodies. Methods Cell Biol 115, 247–267 (2013).

59. M. Mahamdeh, S. Simmert, A. Luchniak, E. Schaffer, J. Howard, Label-free high-speed wide-field imaging of single microtubules using interference reflection microscopy. J Microsc 272, 60–66 (2018).

60. J. Schindelin et al., Fiji: an open-source platform for biological-image analysis. Nat Methods 9, 676-682 (2012).

61. M. Braun et al., Adaptive braking by Ase1 prevents overlapping microtubules from sliding completely apart. Nat Cell Biol 13, 1259–1264 (2011).

62. A. A. Hyman, Preparation of marked microtubules for the assay of the polarity of microtubule-based motors by fluorescence. J Cell Sci Suppl 14, 125–127 (1991).

63. A. D. Cook, S. W. Manka, S. Wang, C. A. Moores, J. Atherton, A microtubule RELION-based pipeline for cryo-EM image processing. J Struct Biol 209, 107402 (2020).

64. S. H. Scheres, RELION: implementation of a Bayesian approach to cryo-EM structure determination. J Struct Biol 180, 519–530 (2012).

65. A. Punjani, J. L. Rubinstein, D. J. Fleet, M. A. Brubaker, cryoSPARC: algorithms for rapid unsupervised cryo-EM structure determination. Nat Methods 14, 290–296 (2017).

66. R. Sanchez-Garcia et al., DeepEMhancer: a deep learning solution for cryo-EM volume post-processing. Commun Biol 4, 874 (2021).

67. P. Emsley, B. Lohkamp, W. G. Scott, K. Cowtan, Features and development of Coot. Acta Crystallogr D Biol Crystallogr 66, 486–501 (2010).

68. E. F. Pettersen et al., UCSF ChimeraX: Structure visualization for researchers, educators, and developers. Protein Sci 30, 70–82 (2021).

69. P. D. Adams et al., PHENIX: a comprehensive Python-based system for macromolecular structure solution. Acta Crystallogr D Biol Crystallogr 66, 213–221 (2010).

70. V. B. Chen et al., MolProbity: all-atom structure validation for macromolecular crystallography. Acta Crystallogr D Biol Crystallogr 66, 12–21 (2010).

