## Supplementary figures and images for "Expanding the Tubulin Code: TTLL11 Polyglutamylase Drives Elongation of Primary Tubulin Chains"

### Supplement Figure 1

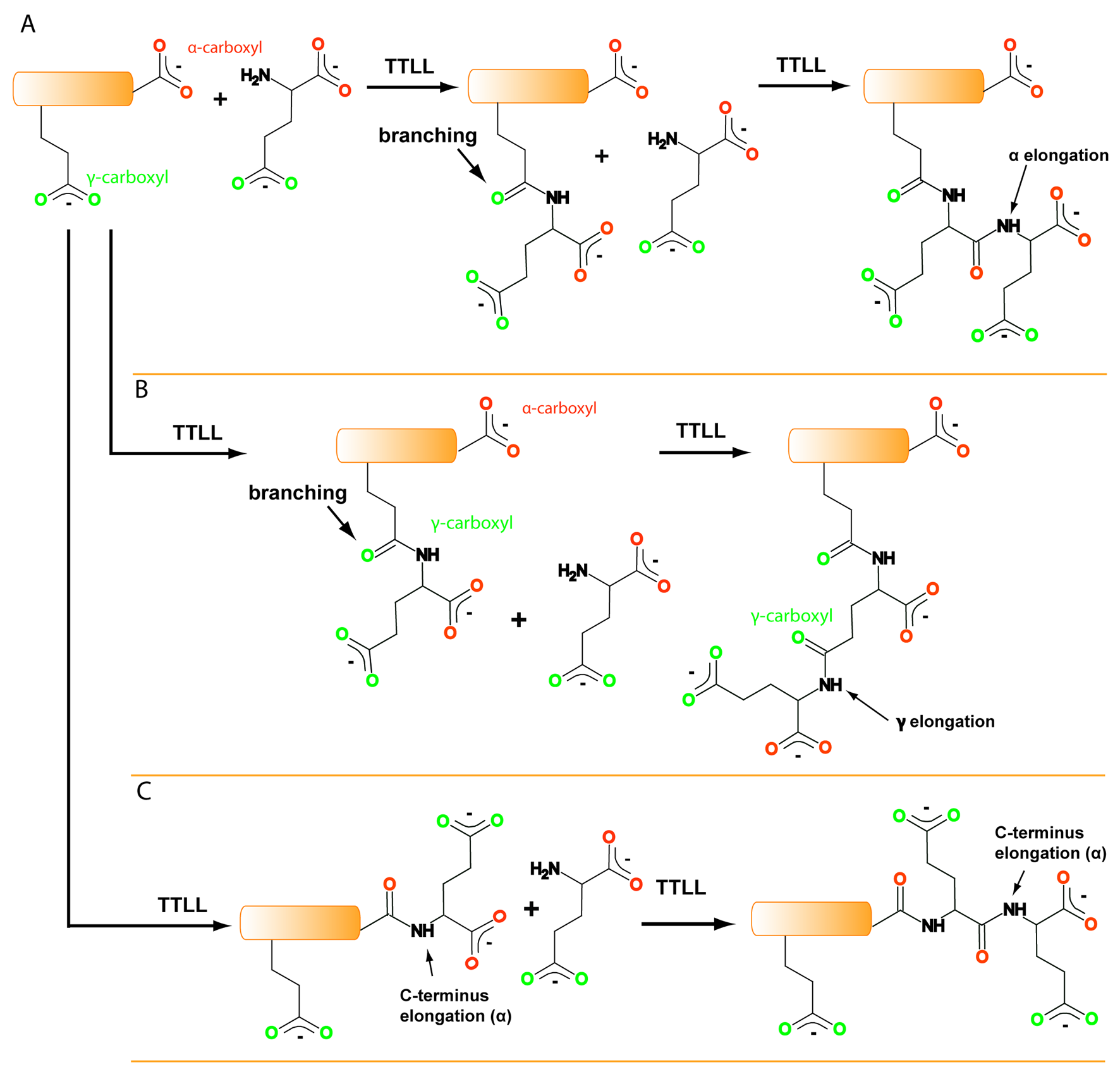

### Supplement Figure 2

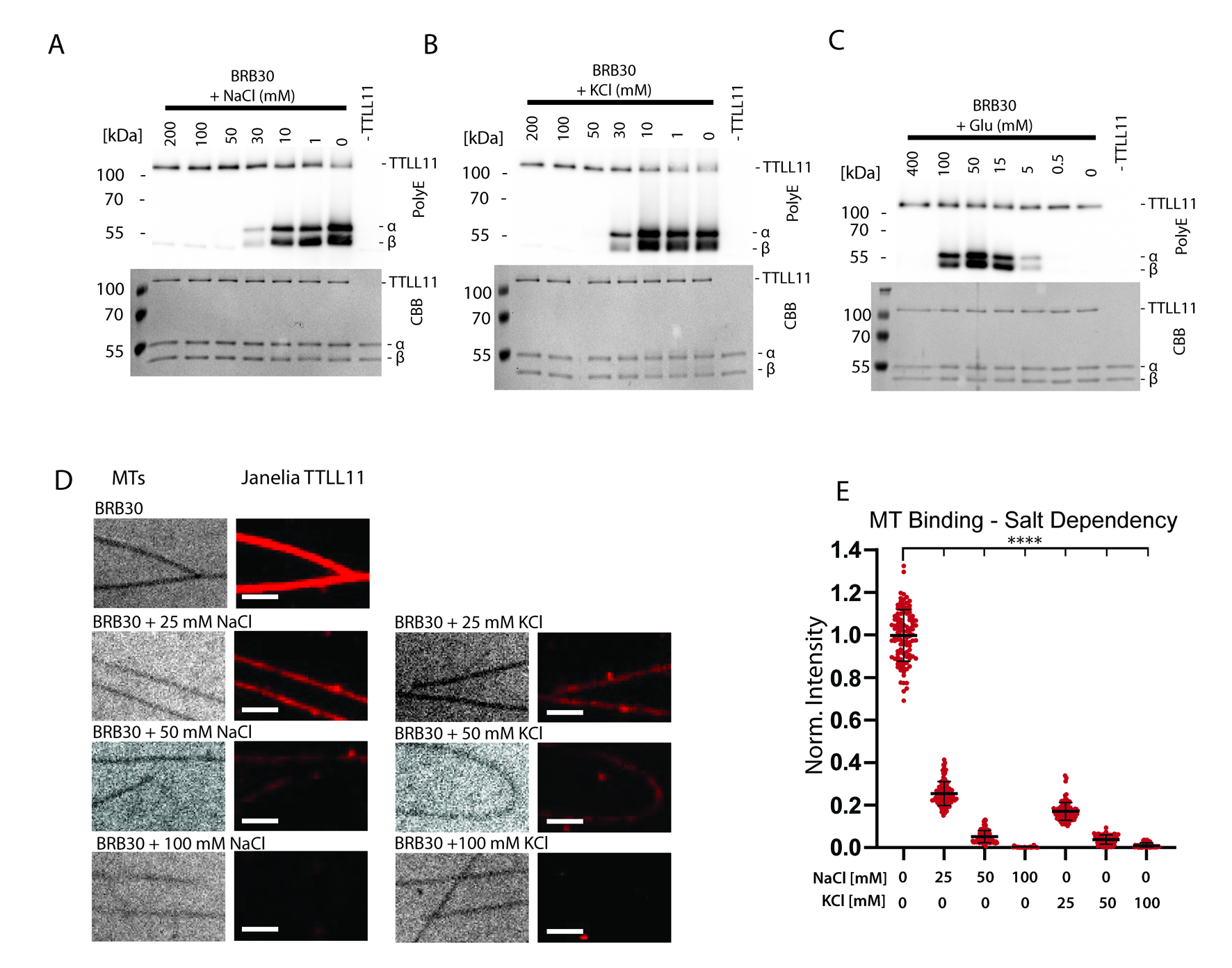

### Supplement Figure 3

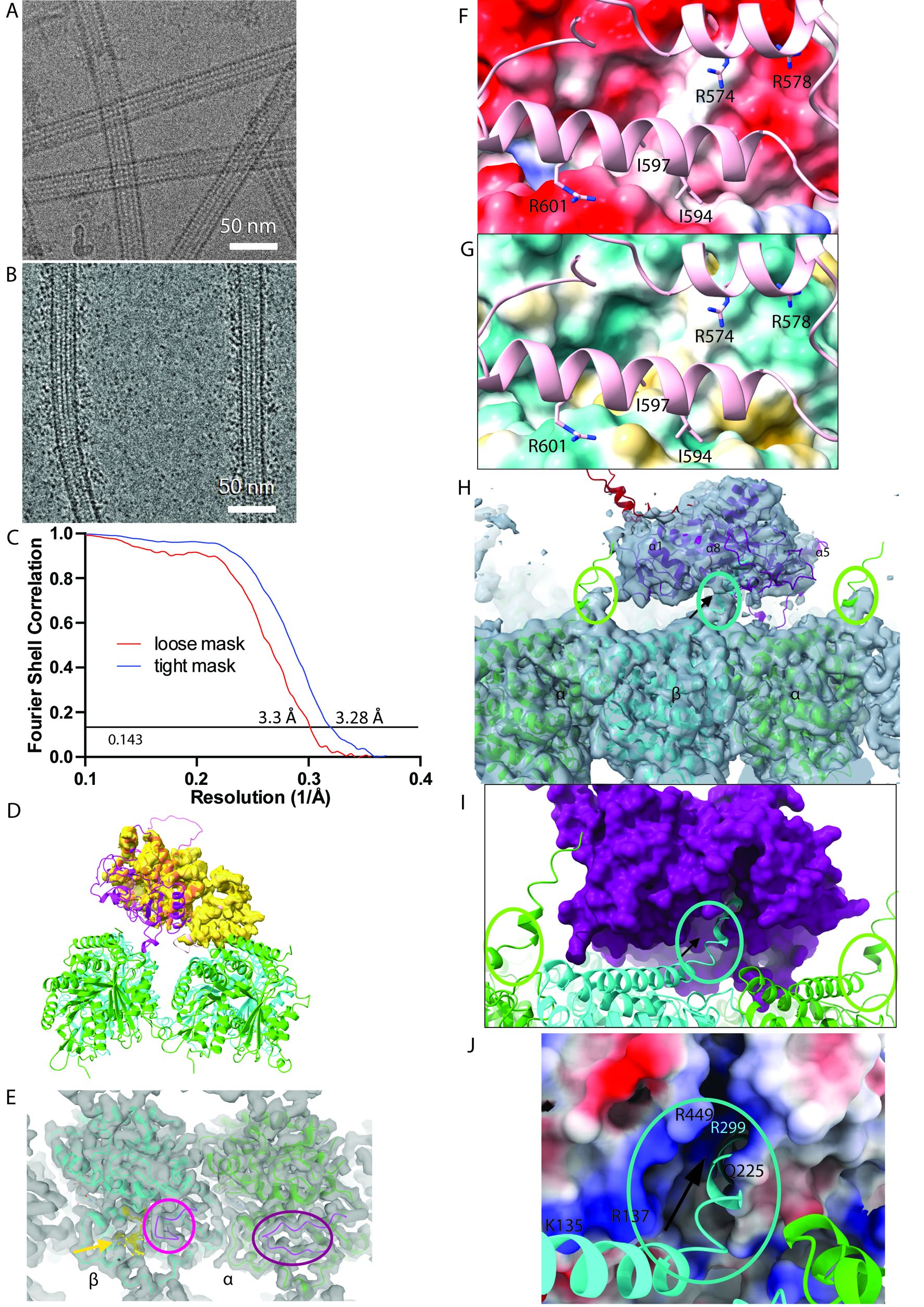

### Supplement Figure 4

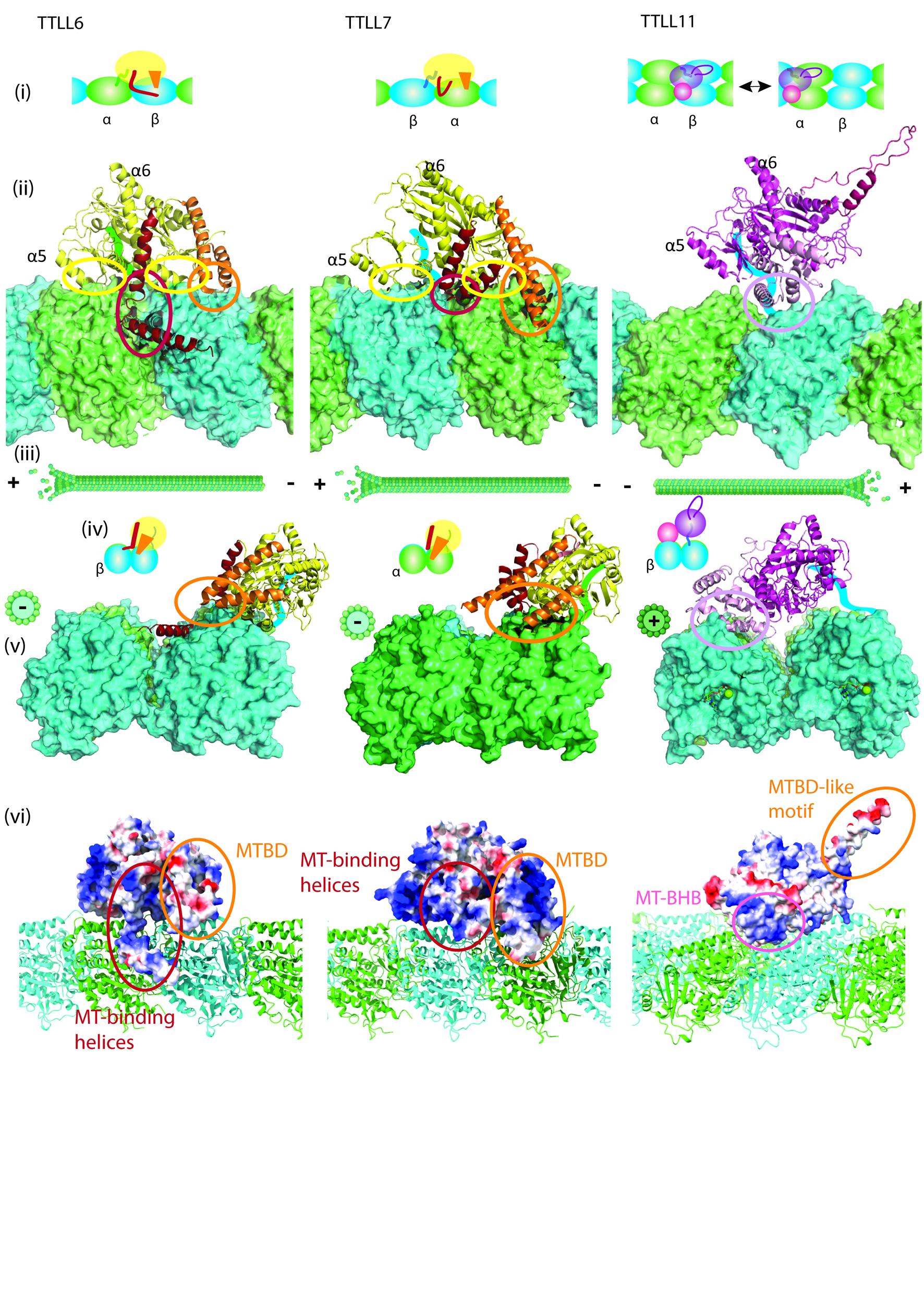

### Supplement Figure 5

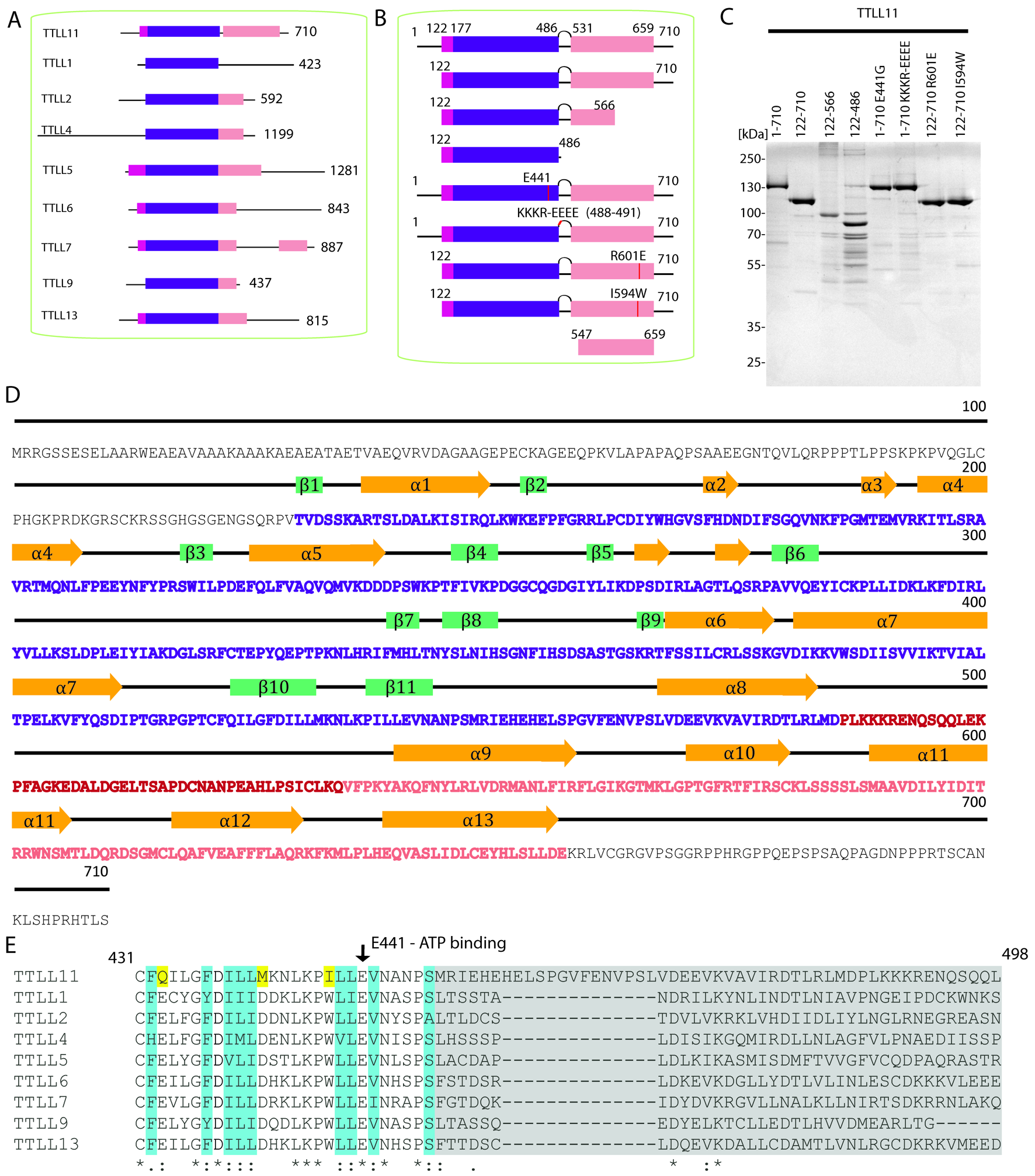

### Supplement Figure 6

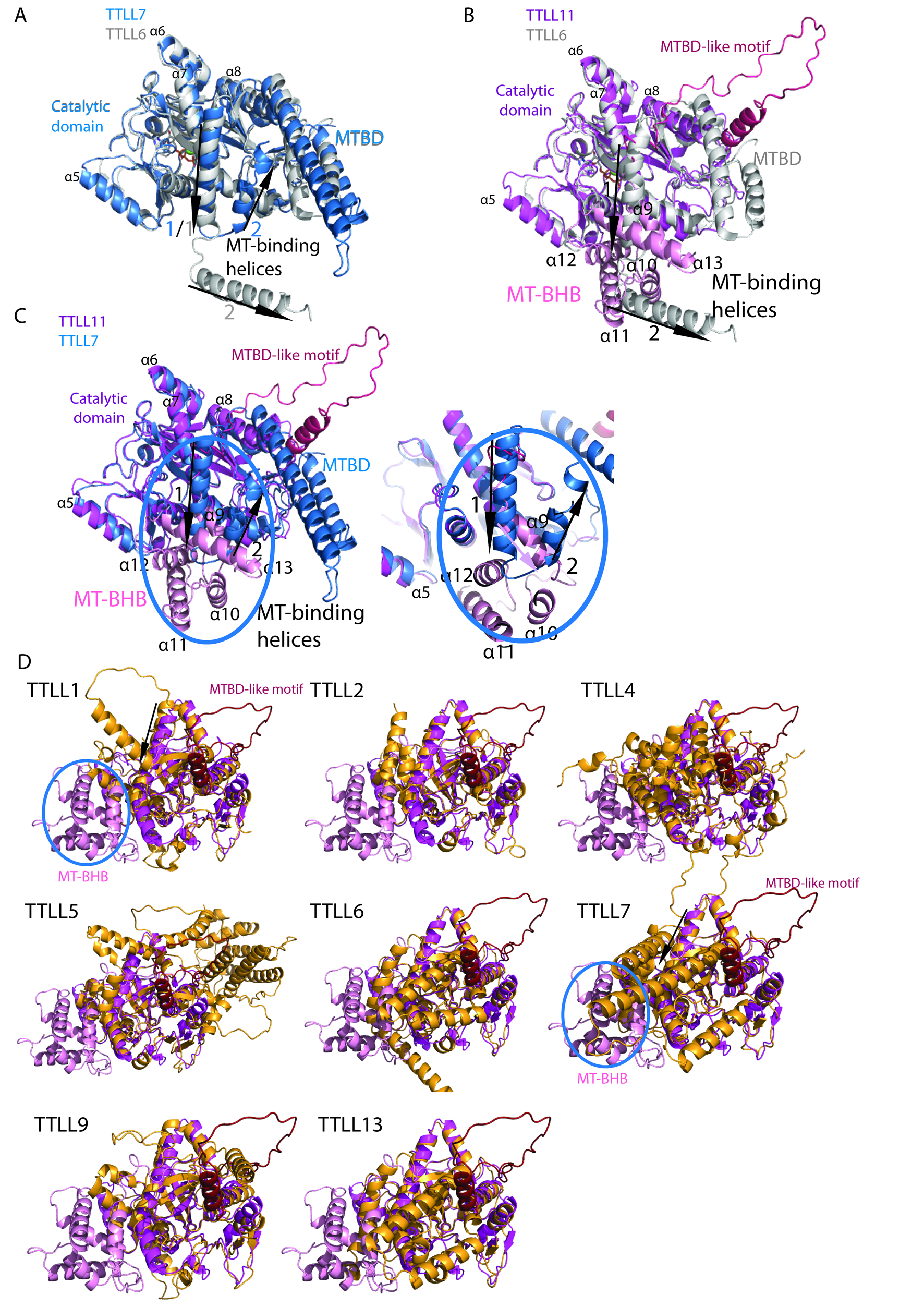

### Supplement Figure 7

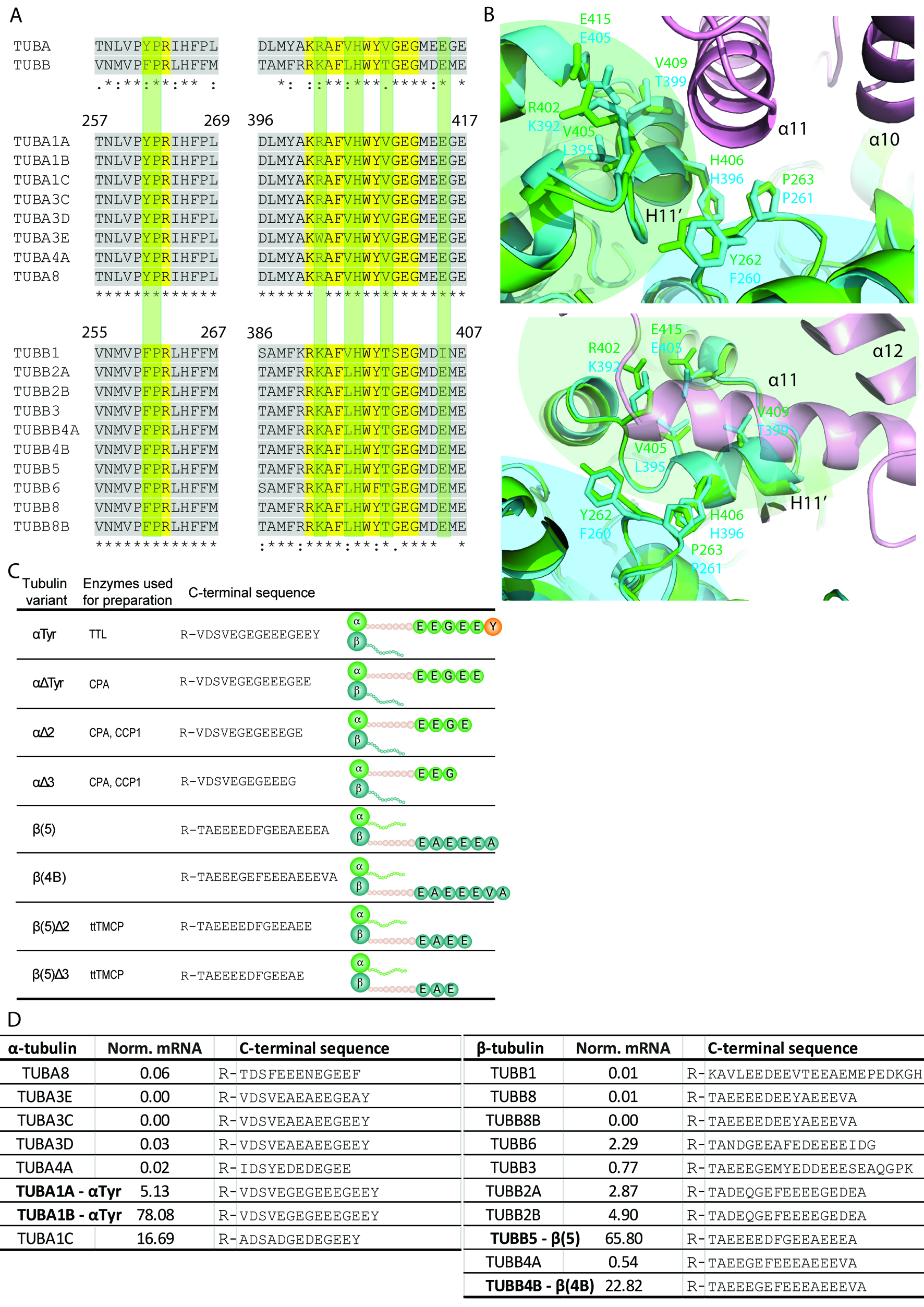

### Supplement Figure 8

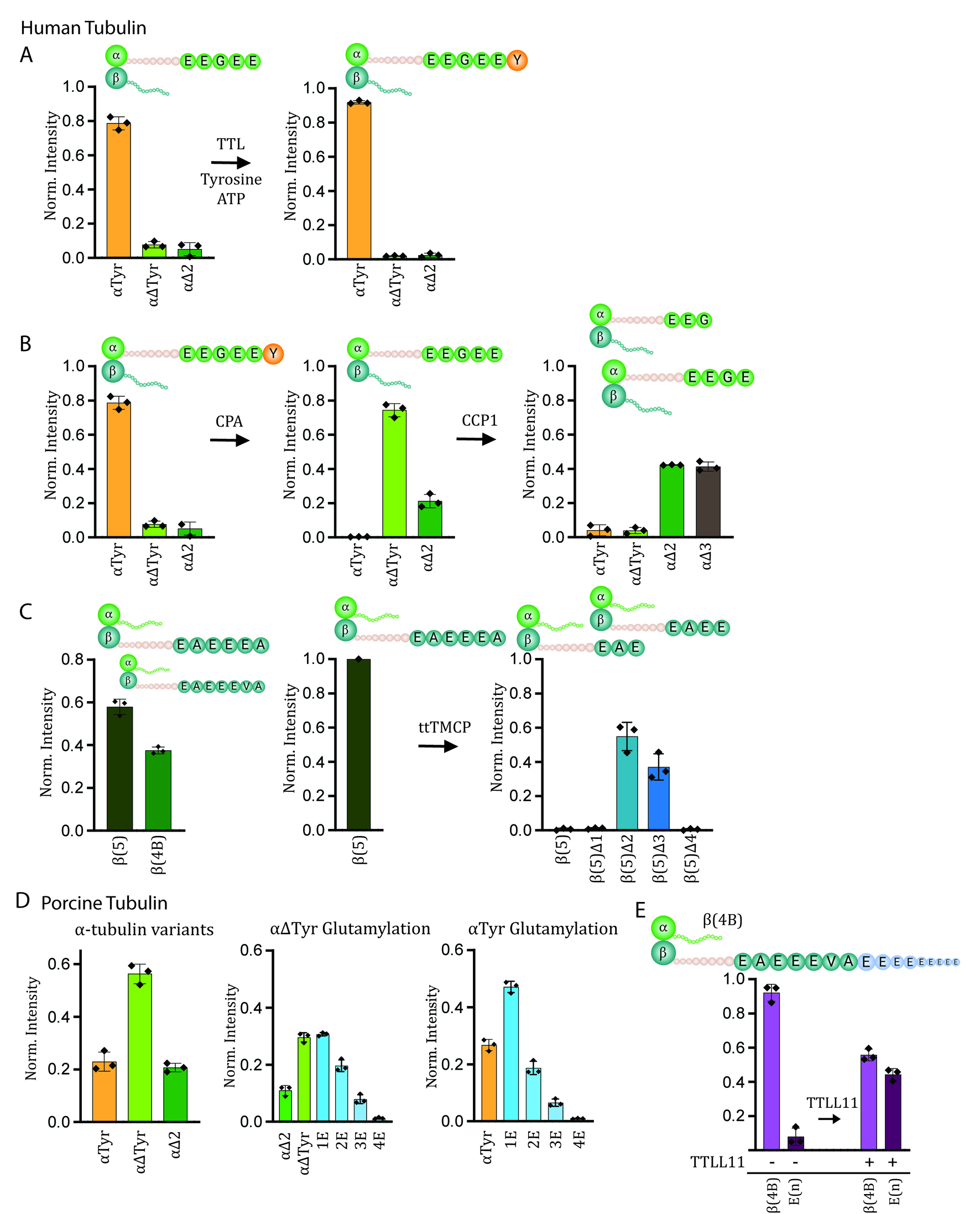

### Supplement Figure 9

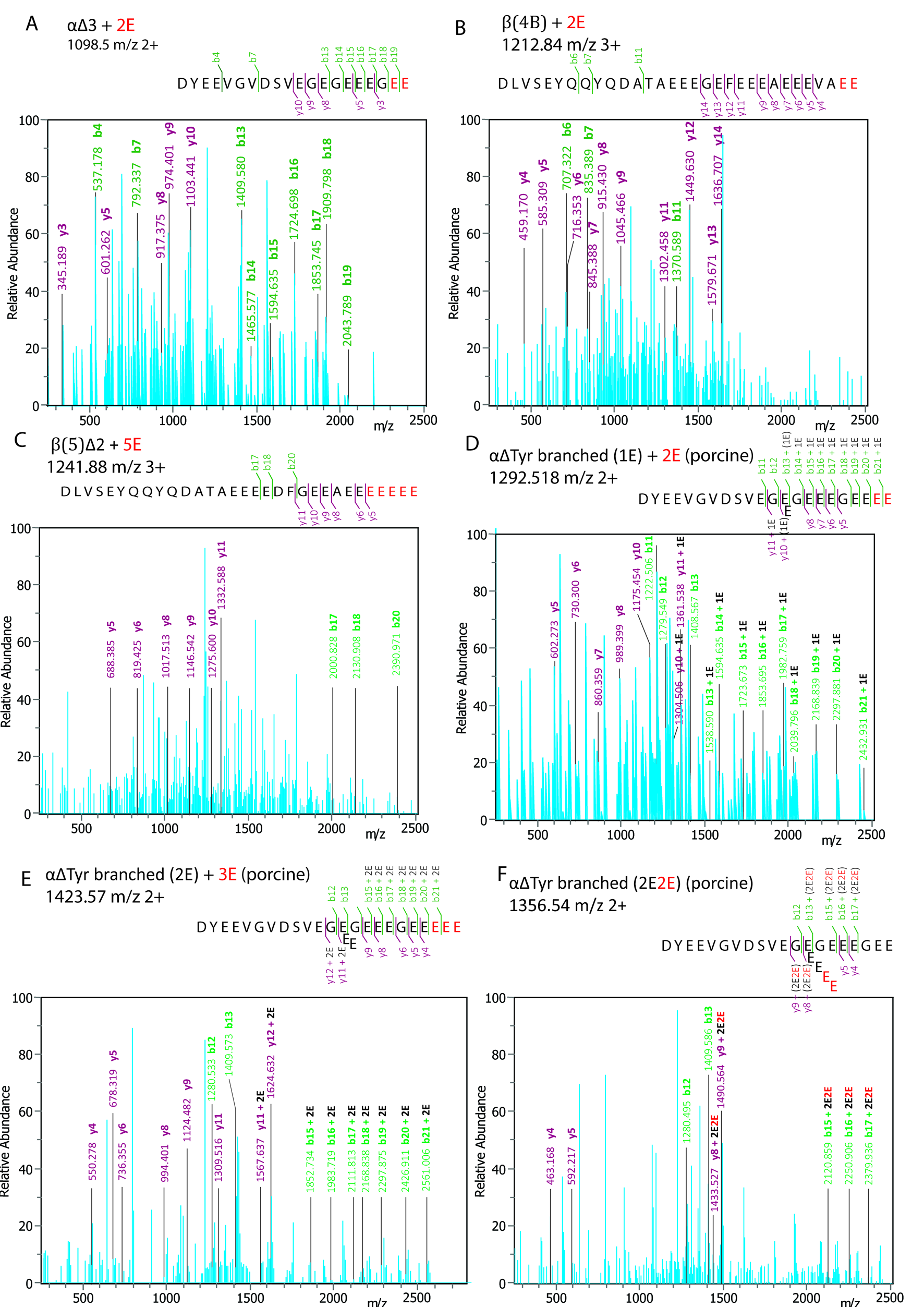

### Supplement Figure 10

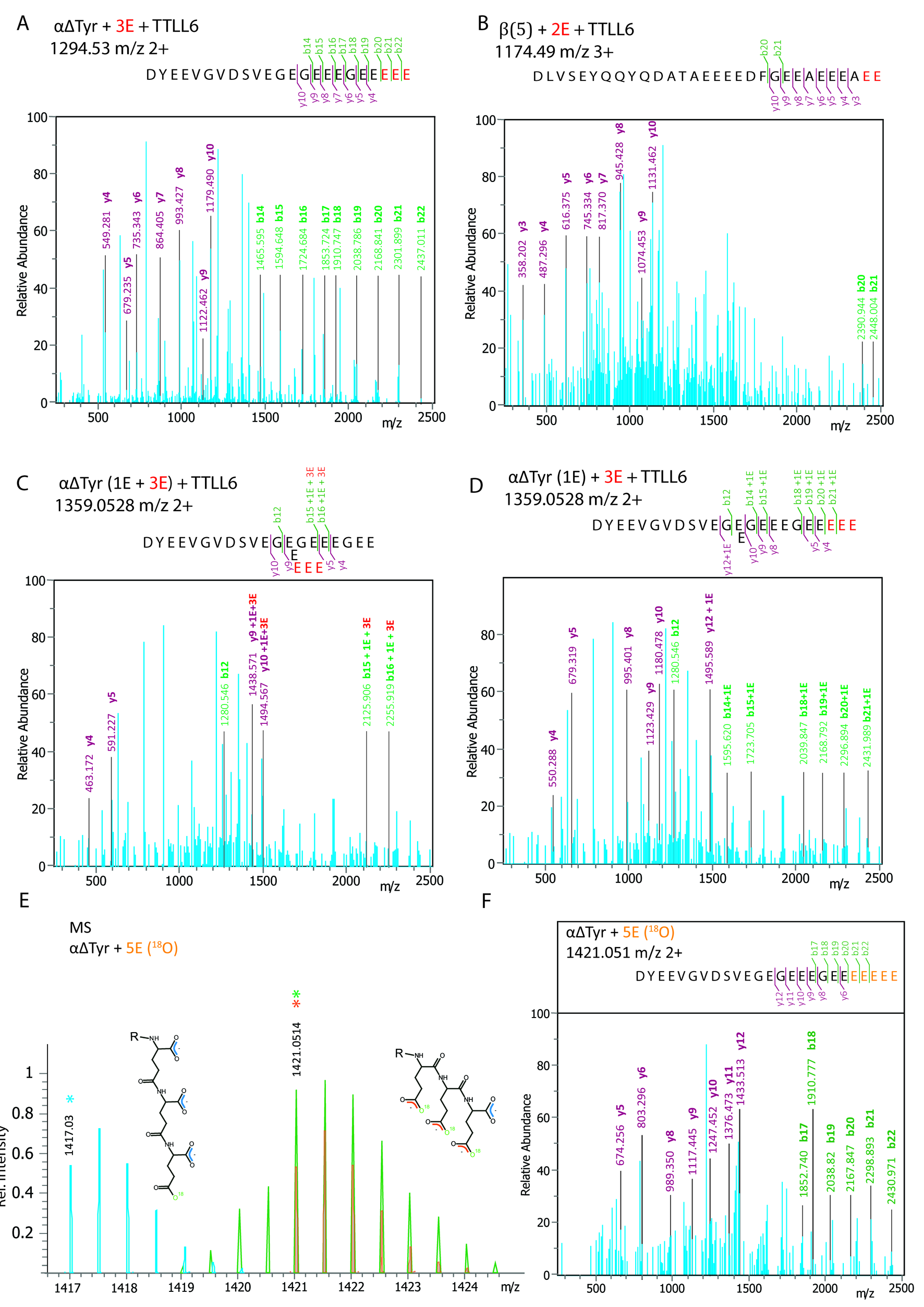

### Supplement Figure 11

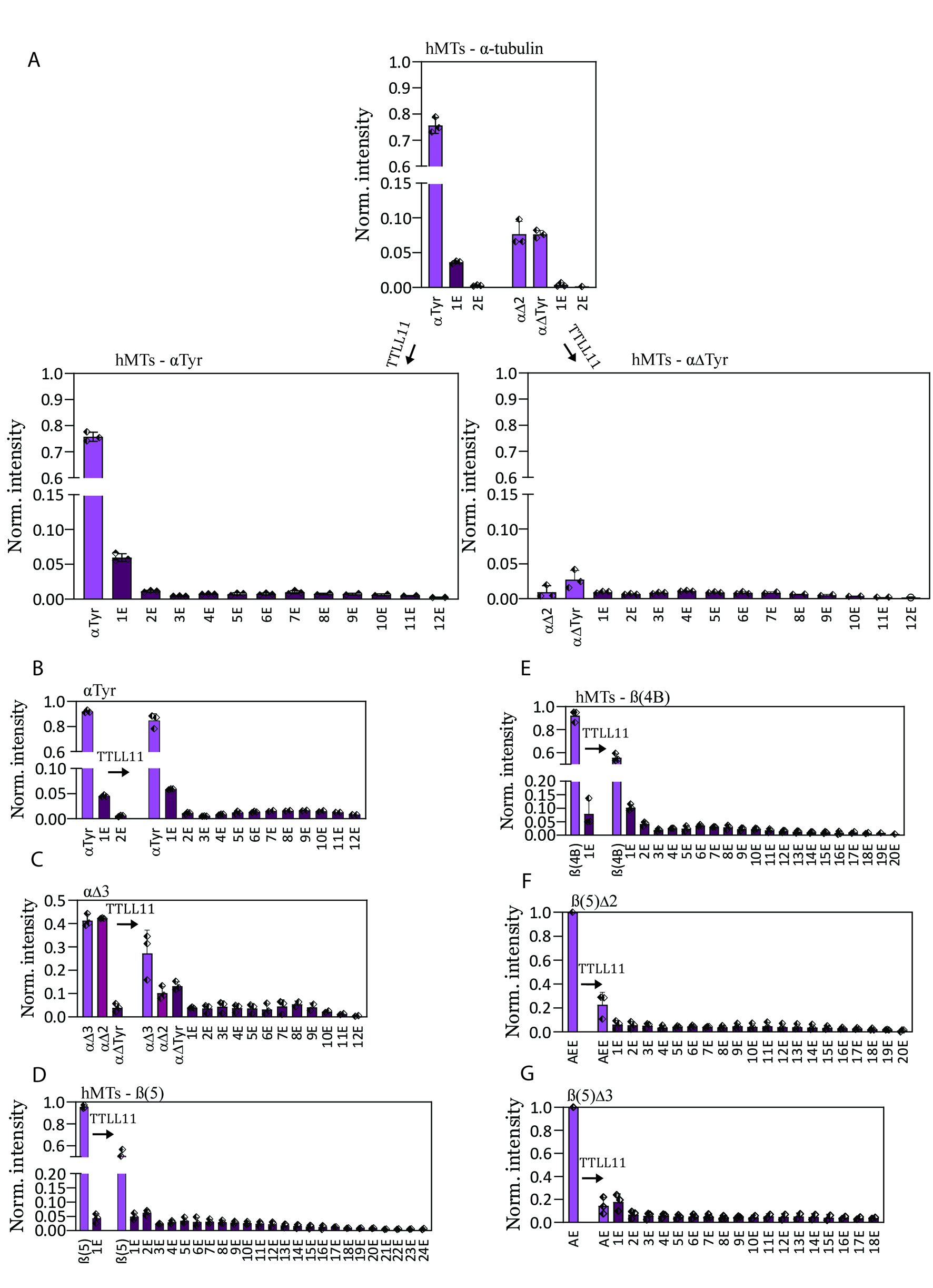

### Supplement Figure 12

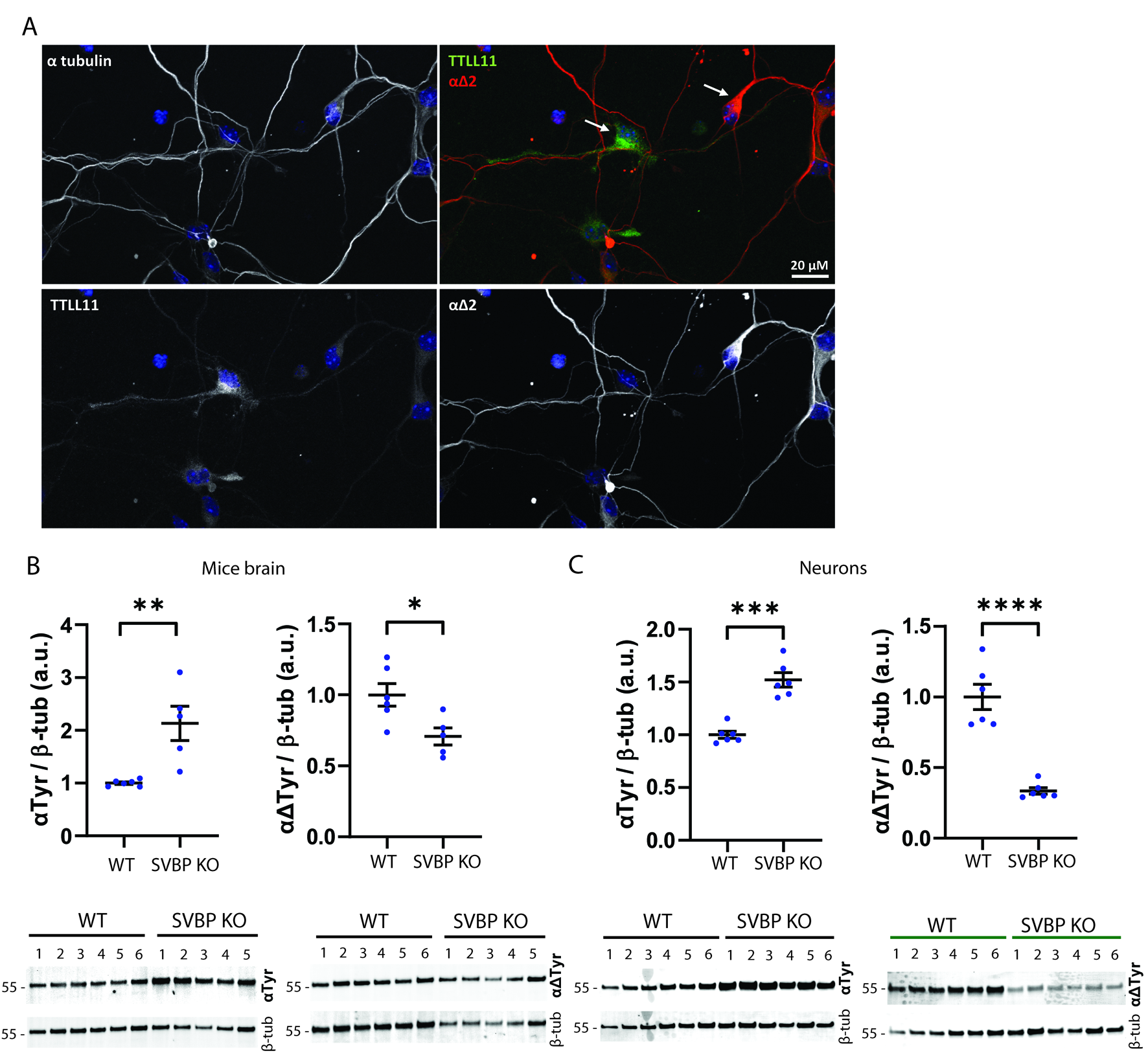
