## Supplement Materials Descrition for "Expanding the Tubulin Code: TTLL11 Polyglutamylase Drives Elongation of Primary Tubulin Chains"

Jana Campbell *et al.*

**Fig. S1. Theoretical polyglutamylation patterns.**

Multiple (theoretical) polyglutamylation patterns can co-exist in a target protein. **A. The current concept of protein polyglutamylation**. During the first initiation (branching) step, a glutamate residue is attached to the γ-carboxyl group of a glutamate side chain within the main peptide chain. The branching point is further extended by adding more glutamates via the α-carboxyl linkage. **B.** **An alternative polyglutamylation pathway.** The initiation step is followed by elongation steps that can attach glutamates via either of α- or γ-carboxylates of the growing polyglutamate chain. **C. A novel polyglutamylation pattern reported here.** The first glutamate residue is attached to the free α-carboxylate of the terminal residue of the main polypeptide chain, resulting in direct extension of the protein sequence. In subsequent steps, the process is repeated as the additional glutamates are attached to the terminal α-carboxyl group.

**Fig. S2. Binding to and polyglutamylation of MTs by TTLL11 - dependency on salt and glutamate concentrations.**

WB showing separation of α- and β-tubulins and detection of polyE chains by the PolyE antibody (upper panel). The corresponding Coomassie–stained SDS-PAGE gel (CBB gel) showing loading of purified proteins (lower panel). TTLL11 activity is negatively correlated with the ionic strength of the assay buffer (**A,B**). TTLL11 activity is the highest between 10 mM - 50 mM glutamate concentration that roughly corresponds to the physiological intracellular glutamate concentration (5 - 20 mM; **C**). **D. Binding of TTLL11 to MT surface depends on salt concentration evaluated by TIRF microscopy.** Microtubules (black) were attached to the glass surface and the binding of TTLL11 (100 nM, conjugated to Janelia-549 fluorophore, red) visualized TIRF microscopy in a buffer comprising 40 mM Tris-HCl, pH7, 5 % glycerol, 1mM TCEP, 1mM MgCl_2_ and varied concentrations of NaCl or KCl (0 – 100 mM). The TTLL11 binding intensity to MTs correlated negatively with an increase of the ionic strength of the assay buffer. Scale bar = 2 µm. **E. Quantification of TIRF images.** The fluorescence intensity of the Janelia-TTLL11 with subtracted background was normalized to MT length and compared the original conditions with added salt concentrations. 50 mM salt decreases the binding almost to the level of background signal. Data is shown as mean fluorescent intensity with n = 2 replicates, with 109, 98, 118, 35, 109, 87, 54 MTs quantified in each sample. Statistical significance was determined using the unpaired t-test with Welsch correction, ****P<0.0001, the black bar represents median value with 95 % c.i..

**Fig. S3. Cryo-EM structure of the TTLL11/MT complex.**

**A,B. Cryo-EM image of MTs and MTs with bound TTLL11** (57 000x magnification), respectively. **C**. **The Fourier shell correlation curves of the cryo-EM map of the TTLL11/MT complex** from Cryosparc with tight and loose mask showing the overall resolution of the map is 3.28 Å. **D**. **The side view of the model of TTLL11/MT complex** with the cryo-EM map representing the TTLL11 protein is shown in gold. Proteins are shown in cartoon representation and colored green, cyan, magenta, and pink for α-tubulin, β-tubulin, the TTLL11 catalytic domain, and the TTLL11 MT-BHB, respectively. **E**. **The cryo-EM map (grey) allows for the unequivocal assignment of α and β-tubulin protomers**. The α-tubulin loop Y357-Q372 (purple oval), and β-tubulin loop D355-K362 (analogous, but 8 amino-acid shorter then in α-tubulin, hot pink oval), and the taxol-binding pocket of β-tubulin (yellow) are highlighted. This difference between the two isotypes is commonly used in cryo-EM to assign the protomers. **F,G**. **Interactions between MT-BHB of TTLL11 and the MT surface**. The α10 and α11 helices of TTLL11 (shown in cartoon representation, pink) represent the primary TTLL11/MT interaction motif. The interaction interface includes (**F**) ionic interactions between the negatively charged MT surface (ChimeraX, colored by Coulombic electrostatic potential red – electronegative, blue – electropositive) and positively charged side chains of R574, R578, and R601; (**G**) hydrophobic interactions between I594 and primarily I597 of TTLL11 and hydrophobic patches at the MT surface (ChimeraX, colored by molecular lipophilic potential green – lipophobic value -20, yellow – lipophilic value 20). **H-J**. **The active site of TTLL11 accommodates the C-tail of β-tubulin**. The TTLL11/MT complex model fitted in the cryo-EM map with a low contour level (0.15). Proteins are shown in cartoon representation and colored green, cyan, magenta/red/pink for α-tubulin, β-tubulin, the TTLL11, respectively. Given their inherent flexibility, only N-terminal segments of tubulin C-tails are visible in the cryo-EM map (green and cyan ovals). Tubulin comprising the “full-length C-tails” were modeled in AlphaFold and manually inserted into the map. The C-terminus of β-tubulin extends into the active site tunnel of TTLL11 (black arrow). **I**. The identical model as in H, with TTLL11 shown in surface representation to better visualize the TTLL11active site tunnel. **J.** TTLL11 shown in surface representation colored by the Coulombic electrostatic potential (ChimeraX, red – electronegative, blue – electropositive) showing the positive charge of the tunnel and its immediate surroundings attracting the negatively charged tubulin tails.

**Fig. S4. Comparison of cryo-EM structures of the TTLL6/7/11 complexes with MTs.**

Structural models and schematic representations of TTLL/MT complexes (front - ii and side view - v) are derived from the structures of TTLL6, TTLL7, and TTLL11/MT complexes (8U3Z (29), provided by Antonina Roll-Mecak (26) and 9HQ4, respectively). Tubulin protomers are shown in surface representation and colored green and cyan for α and β-tubulin, respectively, C-tails that protrude into the active sites of TTLLs are shown as wavy lines of corresponding colors. TTLLs are shown as cartoons and for TTLLs 6/7 the coloring is: the catalytic domain – yellow, MTBD – orange, and the MT binding helices – dark red. For TTLL11: the catalytic domain – magenta, MTBD-like motif – red, the MT-BHB – pink. Interaction interfaces are highlighted by ovals of matching colors. For easier orientation in the positioning of the protein structures, schematic representation of both orientations was added (i, iv). In addition to TTLL interactions with the tubulin C-tails extended to their active sites, TTLLs 6/7 engage MTs by three additional interfaces, while TTLL11/MT interaction interface comprise only helices α10 and α11 of the MT-BHB. The complex interaction interfaces of TTLL6/7/MT, absent in TTLL11, can contribute to the protomer selectivity of the enzymes. The positioning of the catalytic domains in relation to MT differs (iii). In the bottom part (vi), TTLL6/7/11 surfaces are colored by Coulombic electrostatic potential (ChimeraX, red – electronegative, blue – electropositive), tubulins are shown in the cartoon representation. As expected, MT-binding helices as well as MTBD of TTLL7 are positively charged to facilitate interactions with the negatively charged surface of MTs. In the case of TTLL11, MT-BHB is also positively charged while the MTBD-like segment is negatively charged and thus less likely to interact with the MT surface (as also corroborated by our microscopic data).

**Fig. S5. The sequence homology of TTLL glutamylases and TTLL11 variants.**

**A. Schematic representation of TTLL glutamylases.** All enzymes have a high degree of similarity between their catalytic domains (purple), some homologs contain an N-terminal extension (hot pink) tightly interacting thus further being considered a part of the catalytic domain. Lower similarity is observed for N-terminal extension (hot pink) of unknown functions. The C-terminal domains involved in interactions with MTs (MT-binding domains; pink) vary substantially both in length and sequence. Intrinsically disordered N- and C-termini are shown as black lines. **B. Schematic representation of TTLL11 variants used in this study.** The catalytic domain, the N-terminal extension, and the MT-BHB are colored purple, hot pink, and pink, respectively. Intrinsically disordered N- and C-termini are shown as black lines, the MTBD-like motif is shown as the black loop. Site-directed mutants are marked by the red line. **C.** **The CCB-stained gel showing purified TTLL11 variants** comprising the N-terminal TwinStrep-FLAG-HALO tag. Full-length TTLL11 protein (1-710) as well as several truncated variants, designed based on structural predictions and the TTLL11/MT cryo-EM structure, were heterologously expressed in HEK293T cells and purified by the combination of affinity and SEC chromatography. The E531G represents an inactive mutant, K_488_KKR_491_-E_488_EEE_491_ is a mutation of a positively charged motif of MTBD-like motif (originally predicted to be involved in MT binding) (31), and the R691E and I684W are mutations in the MT binding helix α11 of MT-BHB. **D. The sequence and secondary structure prediction of human TTLL11.** The sequence is colored based on the tertiary structure model (black – intrinsically disordered N- and C-terminus, purple – the catalytic domain, dark red – the MTBD-like motif, pink – the MT-BHB) and the secondary structure elements are shown as yellow arrows and green bars for α-helices and β-sheets, respectively. **E.** **A partial sequence alignment of TTLL glutamylases**. The sequences spanning the C-terminal part of the catalytic domain (corresponding to residues C431 – S447) extending into the beginning of MT-binding motives were aligned using Clustal Omega. While there is high sequence conservation within the catalytic domains of all enzymes (blue – similar amino acids, yellow – conserved among TTLLs but different in TTLL11) there is a limited identity sequence beyond the catalytic domain in segments corresponding to the putative MT-binding motives.

**Fig. S6. Superposition of 3D models of TTLL glutamylases.**

**A-C. Superposition of 3D structures of TTLL11/6/7.** 3D structures of glutamylase pairs TTLL6/7 (A), TTLL11/6 (B), and TTLL11/7 (C) were superimposed on corresponding Cα atoms of their catalytic domain (128 – 466 for TTLL11). Structure model of TTLL6, TTLL7, and TTLL11/MT complexes (8U3Z (29), provided by Antonina Roll-Mecak (26) and 9HQ4, respectively).The enzymes are shown in cartoon representation and colored grey, blue, and magenta/hotpink/pink for TTLL6, TTLL7, and TTLL11, respectively. While the structures of catalytic domains are almost identical and superpose well, there are pronounced differences in the structure and positions of putative MT-binding domains/helices. For the TTLL6/7 pair, the helix-loop-helix motifs of the MT-binding domain, which are implicated in MT interactions, are marked by black arrows. The corresponding segment in TTLL11, referred to as the MT-binding domain like motif (MTBD-like), is not visible in the cryo-EM density thus is expected to be flexible and not involved in the MT binding. Instead, the primary TTLL11/MT interface comprises a structurally divergent five-helix bundle (MT-BHB), in which helices α10 and α11 are involved in MT binding. **D. Superposition of 3D models of TTLL glutamylases.** The glutamylases were superimposed on corresponding Cα atoms of their catalytic domains (128 – 466 for TTLL11). The structure of TTLL11 is positioned at 90° left turn from the orientation in A,B,C. Catalytic domain (magenta), MT-BHB (pink, in blue elipse labeled for TTLL1 and TTLL7 alignment), MTBD-like motif (dark red). Superposed TTLLs are in orange cartoon representation. While there is significant structural overlap of the catalytic domains, the MT-BHB is unique for TTLL11 with no structural counterpart in any TTLL polyglutamylase. The predicted internally disordered regiones were deleted for clarity.

**Fig. S7. Tubulin sequence conservation at** **intra- and inter-dimer longitudinal interfaces.**

**A**. Sequences of human tubulin isoforms were aligned using Clustal Omega. Sequences at intra- and interdimer groove interfaces are marked yellow, while residues involved in interactions with TTLL11 are highlighted by green boxes. **B**. **Structural superposition of intra- and interdimer groove interfaces**. 3D structures of tubulin isoforms TUBA1B and TUBB5 (42 % identity, 61 % overall similarity) were superimposed on the corresponding Cα atoms. Given the sequence and structural conservation at both intra- and interdimer groove interfaces TTLL11 cannot effectively discriminate between them. Α and β-tubulin are colored green and cyan, respectively, with amino acids labeled in corresponding colors. The green/cyan background highlight the original tubulin isotype in the cryo-EM structure – upper panel (MT-BHB front view), lower panel (MT-BHB left side view). TTLL11 is colored pink (semitransparent). **C**. **A list of tubulin variants used in this study.** Tubulin isoforms isolated from HEK293T cells primarily comprise isoforms TUBA1A/B, TUBB4B and TUBB5 (our LC-MS data). To enrich individual physiological tubulin variants differing in their C-terminal sequences, which can be underrepresented in HEK293T tubulin, purified tubulins were treated with recombinant tubulin-modifying enzymes (TTL, CPA, CCP1, and ttTMCP) or combinations thereof. **D**. **A list of human α- and β-tubulin isotypes in HEK293T cells** together with normalized mRNA intensities (31) and their corresponding C-terminal sequences. In line with our MS quantification, the most abundant isotypes include TUBA1A/B, TUBB4B and TUBB5. The predominant isoforms that were analyzed by MS in this study are in bold with added acronyms representing the C-terminal sequence.

**Fig. S8. LC-MS based quantification of tubulin variants used in this study.**

**A**. **α-tubulin tyrosination.** αTyr represents the most abundant C-terminal variant of α-tubulins isolated from HEK293T cells, accounting for approximately 80% species, with remains of variants αΔTyr and αΔ2. TTL treatment was used to increase the abundance of the αTyr variant to over 90 %. **B. Generating αΔTyr, αΔ2, and αΔ3 enriched tubulins.** α-tubulins isolated from HEK293 cells contain only approximately 10 % of αΔTyr, while the αΔ3 variant was not detected. CPA treatment was used remove the C-terminal tyrosine to enrich the αΔTyr variant and the subsequent CCP1 treatment to further truncate the C-terminus to enrich the αΔ2 and αΔ3 fractions. **C**. **Generating β(5)Δ2 and β(5)Δ3 enriched tubulins.** β(5) and β(4B) represent the most abundant β-tubulin isotypes isolated from HEK293T cells. As the β(5) isoform is more abundant we focus our attention on this isoform, although both variants were present in reaction mixtures at the same time. β(5) was further modified by the treatment with recombinant TMCP resulting in tubulin fractions enriched in β(5)Δ2 and β(5)Δ3. **D. Tubulins isolated from porcine brains**. Porcine tubulins were isolated by polymerization/depolymerization cycles and individual variants quantitated by LC-MS. Compared to tubulin isolated from HEK293T cells, native porcine brain tubulins are extensively polyglutamylated. **E**. **Polyglutamylation of the β(4B) isoform by TTLL11.** Human MTs were incubated with TTLL11 *in vitro* and glutamylation levels of the β(4B) variant quantified using LC-MS. The glutamylation level upon TTLL11 treatment increased to almost 50 % of all species, the fraction is slightly lower compared to β(5). n = 3, statistical significance was determined using unpaired t-test. The data are shown as the normalized MS intensities, where the sum of intensities of the peptide pool before and after TTLL11 treatment equals 1. *P>0.05.

**Fig. S9.** **The LC-MS/MS analysis of tubulin glutamylation by TTLL11.**

**A-F. MS/MS spectra of polyglutamylated tubulin species.** The LC-MS/MS pipeline was used to analyze tubulin variants that were polyglutamylated by TTLL11 in the presence of isotopically labeled glutamate (D_5_). Peaks of interest observed in b- and y-series of the fragmentation spectrum are highlighted in green and magenta, respectively. This approach enabled the unequivocal assignment of polyE attachment sites within the tubulin sequence. **A**. The αΔ3 peptide with 2 added isotopically labeled glutamates attached directly to the C-terminal glycine. **B**. The β(4B) peptide with 2 added isotopically labeled glutamates attached to the C-terminal alanine. **C**. The β(5)Δ2 peptide with 5 added isotopically labeled glutamates attached to the C-terminal glutamate. **D**. The αΔTyr branched (1 glutamate at E443) peptide from porcine tubulin with 2 added isotopically labeled glutamates attached directly to the C-terminal glutamate. **E**. The αΔTyr branched (2 glutamates at E443) peptide from porcine tubulin with 3 added isotopically labeled glutamates attached directly to the C-terminal glutamate. **F**. The αΔTyr branched (2 glutamates at E443) peptide from porcine tubulin with two added isotopically labeled glutamates attached to the glutamate branch showing this mode of activity is also an option for TTLL11.

**Fig. S10.**  **The LC-MS/MS analysis of tubulin glutamylation by TTLL6 and identification of the linkage of the TTLL11 polyglutamate chains.**

**A-D. MS/MS spectra of polyglutamylated tubulin species.** The LC-MS/MS pipeline was used to analyze tubulin variants that were polyglutamylated by TTLL6 in the presence of isotopically labeled glutamate (D_5_). Peaks of interest observed in b- and y-series of the fragmentation spectrum are highlighted in green and magenta, respectively. This approach enabled the unequivocal assignment of polyE attachment sites within the tubulin sequence. **A**. The αΔTyr peptide with 3 added isotopically labeled glutamates attached to the C-terminal glutamate by TTLL6 showing that this type modification is not limited to only TTLL11. **B**. The β(5) peptide with 2 added isotopically labeled glutamates attached to the C-terminal alanine by TTLL6. **C**,**D**. The αΔTyr peptide branched at E443 was glutamylated by TTLL6 both at the C-terminus (**D**) and the branch (**C**). **E. Theoretical and experimental m/z spectra with ^18^O labeled glutamylation.** MTs were incubated with TTLL11 in the presence of isotopically labeled glutamate ^18^O located at the γ-carboxyl group and the spectrum of the αΔTyr C-terminal peptide extended by five glutamate units analyzed by LC-MS. Theoretical m/z spectra of the αΔTyr-EEEEE peptide with α- and γ-glutamate linkage within the pentaglutamate chain are colored orange and blue, respectively. The experimental m/z spectrum of the peptide (green) overlaps with the orange theoretical spectrum revealing the presence of the α-linkage within the pentaglutamate chain added by TTLL11 as no γ-carboxyl oxygens were released by creation of isopeptide bond. **F**. **The MS/MS fragmentation spectrum of the αΔTyr-EEEEE peptide by ^18^O glutamate.** The LC-MS/MS pipeline was used to analyze tubulin variants that were polyglutamylated by TTLL11 in the presence of isotopically labeled glutamate ^18^O. Peaks of interest observed in b- and y-series of the fragmentation spectrum are highlighted in green and magenta, respectively. The fragmentation spectrum of the αΔTyr-EEEEE peptide reveals that the added glutamates are positioned at the very C-terminus of the peptide (and all are α-linked).

**Fig. S11. The LC-MS quantification of tubulin glutamylation by TTLL11.**

Tubulin variants (in the form of MTs) were incubated with TTLL11 in the presence of isotopically labeled glutamate (D_5_) and intensities of peptides with and without polyE chains attached were summed. The data are shown as the normalized MS intensities, where the sum of intensities of the peptide pool before and after TTLL11 treatment equals 1 (n = 3, technical replicates, statistical significance was determined using unpaired t-test). *P>0.05. **A. Glutamylation of human** **α-tubulins isolated from HEK293T cells.** Upper panel – composition of α-tubulins isolated from HEK293T cells with the majority of the αTyr variant and lesser abundance of αΔTyr/αΔ2 variants. Lower panels – quantification of the individual C-tail peptides with different polyE chains for the αTyr variant (left) and the αΔTyr/αΔ2 variants (right), n = 3. While forming a minority of the tubulin substrate αΔTyr/αΔ2 are preferentially modified by TTLL11 compared to the αTyr variant. **B**. **Glutamylation of the αTyr enriched fractions.** The TTL treatment was used to increase the abundance of the αTyr variant to nearly 100% and then polyglutamylated by TTLL11. In the absence of branching, the αTyr variant is almost not modified by TTLL11. **C**. **Glutamylation of the αΔ3/αΔ2 enriched fractions.** The CPA/CCP1 treatment was used to increase the abundance of the αΔ3/αΔ2 variants that were then polyglutamylated by TTLL11. Both αΔ3/αΔ2 are efficiently glutamylated by TTLL11, although glutamylation of αΔ2 is preferred. **D, E. Glutamylation** **major β-tubulin variants with intact C-termini.** Either of the β(4B) and β(5) variants is efficiently glutamylated by TTLL11 as approximately 50% of the original peptides are glutamylated with up to 20 glutamate units upon TTLL11 treatment. **F, G. Glutamylation of the β(5)Δ2 and β(5)Δ3 enriched fractions.** The β(5) was modified by the treatment with recombinant ttTMCP resulting in tubulin fractions enriched in β(5)Δ2 and β(5)Δ3 variants that were then polyglutamylated by TTLL11. Glutamylation of these variants is very efficient with the majority of the substrate glutamylated in the reaction mixture (around 80 %).

**Fig. S12. αTyr and αΔTyr levels in mice brain and neurons extracts from WT and SVBP KO mice.**

**A**. Confocal images showing a representative example of WT hippocampal neurons transfected with CFP-TTLL11, cultured 2 days *in vitro* and stained for α-tubulin and αΔ2. The CFP signal, amplified using anti-GFP antibody, reveals a cell overexpressing TTLL11 having a lower αΔ2 content in comparison to non-transfected cells. Arrows indicate cell bodies. Scale bar = 20 µm. **B,C**. Immunoblots and their quantification of the ratio of the tubulin modification to β-tubulin in protein samples from 15-weeks old mice brain (**B**) and cortical neurons cultured 8 days *in vitro* (**C**). The absence of SVBP clearly affects tyrosinated/detyrosinated α-tubulin balance in brain and neurons. In neurons, αTyr is increased 1.5-times and αΔTyr is reduced by 70%, similar differences in the brain. Data represent mean ± SEM. n = 6 or 5 animals respectively for WT and SVBP KO, n = 6 independent neuronal differentiation experiment for each genotype. Unpaired t-
