## Supplement Table 1 for "Expanding the Tubulin Code: TTLL11 Polyglutamylase Drives Elongation of Primary Tubulin Chains"

**Table S1 – List of used primers, plasmids and antibodies with their description.**

| **Primers** |  | | | | | |
| --- | --- | --- | --- | --- | --- | --- |
| Primer name | Sequence | | | | | |
| hTTLL11_GTW_1F | GAGAACCTGTACTTCCAGGGCGGAGGCACCATGCGGCGGGGCAGCTCCGAG | | | | | |
| TTLL11 710 GW R | GGGGACCACTTTGTACAAGAAAGCTGGGTTATTA GGACAGGGTATGTCTGGGATGGG | | | | | |
| hTTLL11_GTW_122F | GAGAACCTGTACTTCCAGGGCGGAGGCGAG AACGGCTCCCAGCGGCCGGTC | | | | | |
| hTTLL11_GTW_486R | GGGGACCACTTTGTACAAGAAAGCTGGGTT ATTACTTAAGTGGGTCCATGAGGCGCAG | | | | | |
| mCCP1_1_GW_F | GAGAACCTGTACTTCCAGGGCGGAGGCACC ATGAGCAAGCTAAAAGTGGTG | | | | | |
| mCCP1_1219_GW_R | GGGGACCACTTTGTACAAGAAAGCTGGGTTATTA AATCAGGTGTGTTCTTGATAC | | | | | |
| TTL_1_GW_F | GAGAACCTGTACTTCCAGGGCGGAGGCACCATGTACACCTTCGTGGTACGCG | | | | | |
| TTL_377_GW_R | GGGGACCACTTTGTACAAGAAAGCTGGGTTATTACAGCTTGATGAAGGCAGCTGG | | | | | |
| hTTLL11_R601E_F | CTCTACATTGACATCACA GAG AGGTGGAACTCC | | | | | |
| hTTLL11_R601E_R | GGTCATGGAGTTCCACCT CTC TGTGATGTCAATG | | | | | |
| hTTLL11_I594W_F | GTCCATGGCTGCCG TGG ACTGGCTCTACATTGAC | | | | | |
| hTTLL11_I594W_R | GTGTGATGTCAATGTAGAG CCA GTCCACGGCAGC | | | | | |
| Lenti_expr_1_F | GTCTGATCTCAACAAGCTGTCTAGAGAATTC CGACTCACTATAGGGAGACCCAAGC | | | | | |
| Lenti_expr_R | CAGCTCGACTCAAGAGTCCTCGAGGGCAAACAACAGATGGCTGGCAAC | | | | | |
| Lenti_expr_ins_F | CTCGAGGACTCTTGAGTCGAGCTG | | | | | |
| Lenti_expr_ins_R | GAATTCTCTAGACAGCTTGTTGAGATCAGAC | | | | | |
| Lenti_expr_mid_F | GGTGGATTCCGGTGGTACTGCG | | | | | |
| Lenti_expr_mid_R | CGCAGTACCACCGGAATCCACC | | | | | |
| TTLL11_567_stop_F | CTGGGCATCAAGGGGACAATG taa TTGGGGCCAACAGGCTTTCG | | | | | |
| TTLL11_567_stop_R | CGAAAGCCTGTTGGCCCCAA tta CATTGTCCCCTTGATGCCCAG | | | | | |
| **Plasmids** | |  | |  | | |
| Plasmid name | | Fusion | | Reference | | |
| pEYFP-spacer-hTTLL11_1-710 | | EYFP-hTTLL11 | | Kind gift from Carsten Janke | | |
| pEYFP-spacer-hTTLL11_1-710 E441G | | EYFP-hTTLL11 | | Kind gift from Carsten Janke | | |
| TTL+pWPT | | mTTL | | Marie-Jo Moutin | | |
| p mVASH2 IRES SVBP | | FLAG-VASH2-sfGFP-His-IRES-SVBP | | Marie-Jo Moutin | | |
| pMM322_hTTLL11_1-710 | | Twin-Strap-FLAG-HALO-hTTLL11 | | This study | | |
| pMM322_hTTLL11_122-710 | | Twin-Strap-FLAG-HALO-hTTLL11 | | This study | | |
| pMM322_hTTLL11_122-486 | | Twin-Strap-FLAG-HALO-hTTLL11 | | This study | | |
| pMM322_hTTLL11_212-656 | | Twin-Strap-FLAG-HALO-hTTLL11 | | This study | | |
| pMM322_hTTLL11_547-659 | | Twin-Strap-FLAG-HALO-hTTLL11 | | This study | | |
| pHR_CMV_TetO2_322_mCCP1 | | Twin-Strap-FLAG-HALO-mCCP1 | | This study | | |
| pHR_CMV_TetO2_322_hTTLL11_122-710_E53441G | | Twin-Strap-FLAG-HALO-hTTLL11 | | This study | | |
| pMM322_hTTLL11_122-710_E441G | | Twin-Strap-FLAG-HALO-hTTLL11 | | This study | | |
| pMM322_hTTLL11_122-710_I594W | | Twin-Strap-FLAG-HALO-hTTLL11 | | This study | | |
| pMM322_hTTLL11_122-710_R601E | | Twin-Strap-FLAG-HALO-hTTLL11 | | This study | | |
| pMM322_hTTLL11_122-710_KKKR_EEEE | | Twin-Strap-FLAG-HALO-hTTLL11 | | This study | | |
| pEC566_mTTL_1-377 | | His-MBP-mTTL | | This study | | |
| pMM322 hTTLL6_1-891 | | Twin-Strap-FLAG-HALO-hTTLL6 | | This study | | |
| pMM322 hTTLL7_1-887 | | Twin-Strap-FLAG-HALO-hTTLL7 | | This study | | |
| pMM322 ttTMCP_1-491 | | Twin-Strap-FLAG-HALO-ttTMCP | | This study | | |
| **Antibodies** | |  |  | |  |  |
| Antibody name | | Source organism | Company | | Dilution | RRID |
| anti-Polyglutamate chain (polyE) | | Rabbit | AdiopoGen | | 1:4000 | AB_2490540 |
| Anti-alpha Tubulin antibody - Microtubule Marker | | Rabbit | Abcam | | 1:2000 | AB_2210057 |
| Anti-β-Tubulin antibody | | Mouse | Sigma | | 1:2000 | AB_477556 |
| Glu-alpha-Tubulin (Detyrosinated alpha-Tubulin) (alpha) Rabbit Polyclonal Antibody | | Rabbit | OriGene AP54970SU-N | | 1:2000 | - |
| Anti-α-Tubulin antibody, tyrosinated, clone YL1/2 | | Rat | Sigma | | 1:2000 | AB_2890657 |
| anti-polyglutamylation Modification, mAb (GT335) | | Mouse | AdipoGen | | 1:4000 | AB_2490211 |
| Precision Protein StrepTactin-HRP Conjugate | | StrepTactin-HRP | BioRad | | 1:4000 | - |
| Anti-Rabbit IgG - Peroxidase antibody produced in goat | | Goat | Sigma | | 1:10000 | AB_258284 |
| Anti-Mouse IgG - Peroxidase antibody produced in goat | | Goat | Sigma | | 1:10000 | AB_258167 |
| tubulin α ∆2 | | Rabbit | Moutin Lab | | 1:2000 | - |
