## Supplement Table 2 for "Expanding the Tubulin Code: TTLL11 Polyglutamylase Drives Elongation of Primary Tubulin Chains"

**Table S2 - Cryo-EM data collection, model refinement and validation statistics.**

| **Data collection** |  |
| --- | --- |
| Microscope | Titan Krios (G1-2) |
| Detector | Falcon 3EC direct electron camera |
| Voltage (kV) | 300 |
| Defocus range | -0.8 - -2 μm |
| Pixel size | 1.349 Å |
| Frames per movie | 30 |
| Movies collected | 8.8k |
| **Image processing** |  |
| Symmetry | C1 |
| Initial particle images | 250k MT segments |
| Final particles images | 240k subtracted segments |
| Map resolution | 3.2 Å |
| FSC threshold | 0.143 |
| Map resolution range | 3 to 11 Å |
| **Model refinement and validation statistics** |  |
| Atomic modeling refinements packages | Coot, Phenix |
| Initial model used | AF-Q8NHH1-F1-v4 |
| Model Resolution (Å) | 3.2 |
| FSC threshold | 0.143 |
| **Model composition** |  |
| Non-hydrogen atoms | 17592 |
| Protein residues | 2230 |
| **Ligands** | 4 GMPCPP, 4 Mg^2+^ |
| **B factor (Å^2^)** | 102.3 |
